# Tubulin C-terminal tails are pH sensors that regulate microtubule function

**DOI:** 10.64898/2026.03.06.710195

**Authors:** A.M. Whited, Patrick DeLear, Ezekiel C. Thomas, Jeffre Allen, Génesis Ferrer-Imbert, Nirbhik Acharya, Carlos A. Castañeda, David Sept, Jeffrey K. Moore, Loren E. Hough

## Abstract

Changes in intracellular pH are critical for maintaining homeostasis, mediating signaling pathways, and enabling cellular responses to stress, injury, and disease. There is increasing evidence that clusters of acidic residues, primarily glutamates, are both highly prevalent and conserved in disordered regions of proteins and can play an important role in cellular pH response. Tubulin C-terminal tails (CTTs) are glutamate rich regions which protrude from the microtubule surface. These tails are a primary site of for both post-translational modifications and binding of microtubule-associated proteins. Motivated by these observations, we measured the pH response of tubulin CTTs using NMR spectroscopy, circular dichroism, and computational simulations. We find that glutamate residues in CTTs taken from organisms across eukaryotes exhibit a robust upshift in their p*K*_a_ values, that the sequential context of glutamate residues creates hot spots for protonation, and that hydrogen bonding between side chains stabilizes interactions that alter the conformation of the CTT. To determine whether the CTT pH response plays a potentially important role in microtubule interactions, we measured the pH dependence of the binding of the yeast kinesin-5, Cin8, to microtubules. We find that Cin8 binding is modulated by pH in a CTT-dependent manner. Our results demonstrate that acidic clusters are important mediators of cellular pH response and establish that pH can regulate interactions at the microtubule surface.

**Significance Statement:** Variation in cellular pH is important for cell function in changing environmental conditions or developmental states. Here we probe protonation of the glutamate-rich C-terminal tails of tubulin, revealing the existence of and mechanism driving the anomalously high pH response and subsequent regulation of microtubule binding. Our results demonstrate that acidic clusters are important mediators of cellular pH response and establish pH-based regulation of interactions at the microtubule surface.

## Introduction

While tightly regulated, the cytoplasmic pH (pH_IN_) shows significant fluctuations as a function of cellular state, including during the cell cycle, development, in injury or disease, and as a function of organism or tissue type (1–9). While pH_IN_ is typically assumed to be around 7.2, observations of mammalian and yeast cells pH_IN_ range from 5.3 and 7.6 (1– 5, 10–13). There is growing evidence that a wide range of cellular processes respond to changes in pH_IN_. Acidification of pH_IN_ during ischemia is thought to be strongly protective of cardiac tissue through regulation of systems as diverse as contractile machinery, the mitochondria and proteases (14). The atypical regulation of intracellular pH in cancer affects a range of cellular processes, including glucose metabolism, migration and cell cycle progression (4, 15, 16). Moreover, changes in pH need not be large to direct signaling processes in mammalian cells, with cellular responses seen for pH shifts as small as 0.2 to 0.3 units (3, 17).

The most well understood molecular mechanisms of cellular pH responses typically occur through changes in the protonation state of ionizable residues in ordered structures, where the local chemical environment is tuned to shift the p*K*_a_ of amino acid side chains to the relevant physiological range. This is most well studied in enzyme active sites, including many examples of pH regulation in the cytoskeleton (18–27). Increasingly, protonation resulting from the physiological variation in pH_IN_ is being described as a form of post-translational modification (PTM), because protonation regulates protein function though altering enzymatic activity or binding interactions (28–30). This effect is akin to traditional PTMs, though the protonation state is regulated by global pH_IN_ changes, rather than by specific modifying enzymes.

While histidine protonation is commonly seen as the primary mechanism of pH regulation in disordered regions (31–33), there is increasing evidence that residues within acidic clusters, particularly glutamates, can be protonated in physiological conditions. Acidic residues typically have p*K*_a_ values around 4, far below the typical physiological pH range (20). Nonetheless, glutamate protonation is implicated in the pH gradients observed in the nucleolus, in regulation of G3BP1/2 condensation, in the stress- and pH-dependent phase separation of Sup35 in yeast, and in the aggregation of prothymosin α (34–38). Moreover, clustering of glutamate residues in disordered regions is a prevalent and conserved feature in disordered proteins, suggesting their functional importance (39).

However, it remains to be determined the extent to which the pH-sensing of glutamate-rich regions is a general feature. The molecular mechanisms underlying the physiological pH response from an amino acid with a relatively low p*K*_a_ value are not well understood, nor are the sequence features that enable such a pH response, or how or whether the observed pH response elicits a biological outcome. Given the potential for widespread use of glutamate clusters in pH sensing, we sought to determine whether pH sensing was a conserved feature of glutamate clusters in an orthogonal biological system to those already considered(34–36).

In nearly all eukaryotes, α- and β-tubulins contain glutamate-rich C-terminal tails (CTTs). Many eukaryotes express multiple genes encoding for α- and β-tubulins, known as isotypes. The result is a rich library of protein sequences to probe glutamate-mediated pH responses (40). Microtubules, built from α- and β-tubulins, play a central role in cellular signaling, mobility and division, all cellular functions regulated by changes in pH_IN_ (28, 41). The CTTs protrude from the microtubule surface and are therefore a primary site of binding of microtubule-associated proteins (MAPs) and of PTMs (42). Knowing that microtubules play a central role in changes in cell behavior broadly, we sought to determine whether and how CTTs respond to changes in pH, and whether these responses were important for the regulation of microtubule function.

Here we investigate the pH sensitivity of tubulin CTTs and the effects of protonation on the CTT conformational ensemble and binding. We use a combination of nuclear magnetic resonance (NMR) spectroscopy, circular dichroism (CD), molecular dynamics simulations, and reconstitution experiments to measure pH-dependent protonation, protein conformational changes and protein binding. The measured p*K*_a_ values in the CTTs studied are shifted significantly higher than from their model values (20). We identify two primary mechanisms driving this shift: 1) electrostatic repulsion within clusters of glutamate and aspartate residues, 2) hydrogen bonding with nearby glutamate side chains which stabilizes a residue in its protonated state. Hydrogen bonding results in sustained changes in the conformation of the CTT. We find that protonation regulates the binding of kinesin-5/Cin8 to microtubules and so has the potential to play an important role in microtubule function. Collectively, our findings establish tubulin CTTs as pH sensors for regulating microtubule function in cells. We propose a model where the syntax of polyglutamate-rich motifs potentiate protonation events, offering a new rationale for the evolution of glutamate patterning in tubulin CTTs. Given the prevalence of glutamate-rich clusters in intrinsically disordered regions (IDRs) (43), our findings provide insight in how disordered regions in proteins sense changes in pH that drive cellular function.

## Results

### TUBA1A^CTT^ shows an anomalously high pH_m_ and strong pH response

To test the possible role of glutamate-rich CTTs in the pH response of tubulin, we purified the CTT peptide of TUBA1A (TUBA1A^CTT^, Fig. 1A, S1) for subsequent studies by both CD and NMR. Our aim was to probe for changes in the structural composition of the conformational ensemble of the TUBA1A^CTT^ as pH is changed. While our interest lies primarily in the physiological pH range, we performed measurements over a broad pH range to better quantify the pH response.

**Figure 1.**
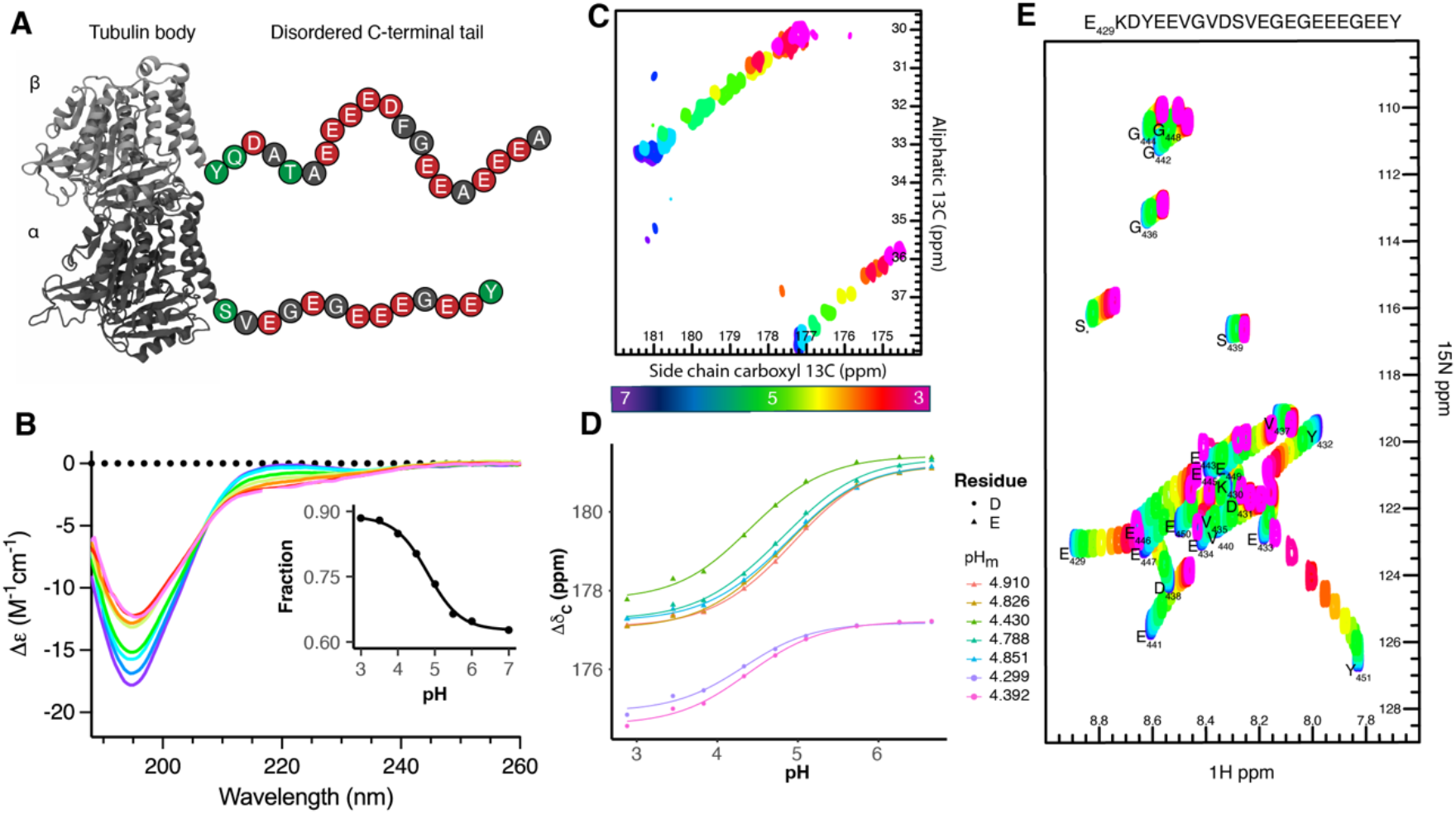
Circular dichroism and NMR show significant response of the TUBA1A^CTT^ to changes in pH. A) Cartoon representation of the tubulin dimer highlighting the prevalence of acidic residues. B) Circular dichroism spectra of TUBA1A^CTT^ as a function of pH (pH 7(purple), 6, 5.5, 5, 4.5, 4, 3.5, and 3 (red)). The inset shows the magnitude of the component of the SESCA deconvolution using the DS-B4R1 basis set. C) CBCGCO experiment showing the chemical shift changes as a function of pH of the glutamate and aspartate side chains. D) Quantification of the side chain carbonyl chemical shifts as a function of pH, fit with the Henderson-Hasselbalch equation, with half maximum of the pH response listed in the legend. E) ^15^N HSQC spectra showing pH of response backbone amides for each residue.

The CD spectra of the same TUBA1A^CTT^ peptide show that the peptide remains primarily disordered between pH 7 and 3, though with some change in the conformational ensemble, as revealed by changes in the depth of the well at 195 nm, and the broad shoulder at around 220 nm (Fig. 1B). The observed spectra are consistent with a pH-dependent change in polyproline II conformational propensity (44, 45). To quantify the spectral changes as a function of pH, we deconvolved the CD spectra using SESCA and a basis set (DS-B4R1) which has been optimized for disordered proteins (46)(Fig. 1B). The fit pH_m_, or the half maximum of the pH response, of the curve was 4.8 ± 0.1, significantly higher than the p*K*_a_ value of isolated glutamate residues (4.2 ± 0.1) in disordered loops (20). If we take the overly naive (as discussed below) assumption that the acidic residues in TUBA1A^CTT^ respond independently to pH, then the probability of TUBA1A^CTT^ being protonated on at least one of the acidic groups is 15% at pH 6.5, a pH relevant for ischemia and cancer (Fig. S2) (47–49).

To further assess the effects of protonation on individual amino acids within the TUBA1A^CTT^, we performed NMR-based pH titration experiments between pH 7 and 3 and again found unusually high pH_m_ values (Fig. 1C-E). We first measured the chemical shifts of the side-chain carbon atoms nearest the titratable oxygen (C_γ_/C_δ_ and CO). The pH_m_ values measured centered on 4.8 for the glutamate residues, and 4.3 for the aspartate residues, which are ∼0.5 pH units higher than Glu and Asp model compound p*K*_a_ values, respectively (20). The resonances of the same amino acid type overlapped throughout the pH titration, indicative of highly similar chemical microenvironments for the side chains of these residues in the sequences. As a result of this overlap, we could not reliably distinguish the individual titration curves of different residues.

To probe for differences of the pH response among different residues, we then turned to ^1^H-^15^N HSQC experiments that probe perturbations to the backbone amide chemical shifts (Fig. 1E). Consistent with widespread observations that the CTTs are fully disordered (50, 51), the NMR HSQC spectra show ^1^H chemical shift values clustered between 7.8-8.8 ppm, a region characteristic of the fast-tumbling times of disordered regions (52). Although the chemical shift changes of the amide resonances are sensitive to changes in structural or bonding changes, they can be useful monitors for nearby side chain protonation events (53).

The HSQC spectra showed large, coordinated chemical shift perturbations for most amino acids of TUBA1A^CTT^, again with a higher than typical apparent pH_m_ (20)(Fig. 1E). The magnitude of the amide chemical shift changes are consistent with the formation of intra-molecular hydrogen bonds between protonated glutamate side chains and backbone amide residues (21, 54). Large chemical shift changes can also be due to significant changes in secondary structure, although our CD data excludes this possibility.

### Pairwise residue interactions are described by principal component analysis

Because of the coordinated response across the peptide, we applied principal component analysis (PCA) to our HSQC pH titration data (Fig. 2A, S3) (55). As protonation events are significantly faster than the timescales probed by NMR, we hypothesized that PCA would reveal the core features of the pH response. We found that over 99.7% of the pH-dependent changes to the TUBA1A^CTT^ NMR spectra is explained by the first two principal components.

**Figure 2.**
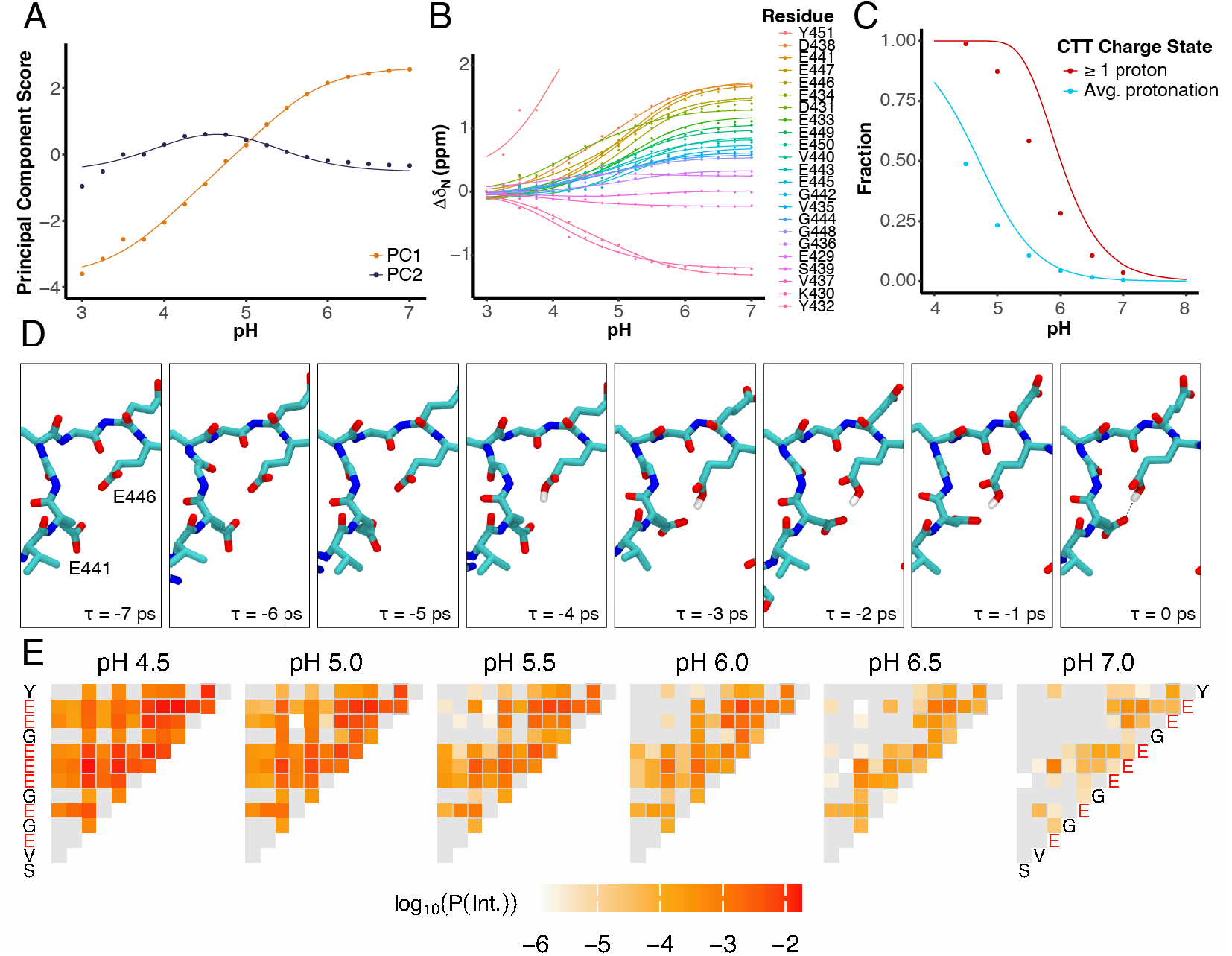
Interactions between residues drive a strong pH response. A) The pH response of the first two principal components with fits to Eq. 1, showing that the response can be described using two pseudo-site p*K*_a_ values. B) The PCA fits were mapped back to the individual residues, showing strong agreement as seen here for the ^15^N amide chemical shifts as a function of pH. C) Simulation results showing both the average protonation fraction and the fraction of time each peptide had at least one protonated acidic residue. Points are from simulations, while curves assume an average p*K*_a_ value taken from the carbonyl side chain experimental data (Fig.1D). D) Example simulation trajectory resulting in protonation and subsequent hydrogen bonding stabilizing a bent conformation of the peptide. E) Heat maps showing the prevalence of pairwise interactions resulting from protonation.

Interactions between residues could explain the upshift in p*K*_a_ and would be expected to be apparent in our PCA data. It has been well established that a system of two *interacting* charged residues have a pH response that is a linear combination of the pH response of two *isolated* charges with different p*K*_a_ values (56, 57). Indeed, both the first and second principal component scores are well described by a two-site model with two pseudo p*K*_a_ values, p*K*a_1,2_. The first two principal components are well fit by the sum (PCA_1_) and difference (PCA_2_) of the typical Henderson-Hasselbalch response of two independent charges with these two pKa values:

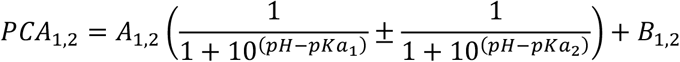

where A_1,2_ and B_1,2_ are scaling constants and p*K*a_1,2_ are pseudo p*K*_a_ values. The fitted pseudo p*K*_a_ values are 4.0 ± 0.2 and 5.2 ± 0.1, consistent with our model of two interacting ionizable groups. Having fit the PCA scores, we then used the coefficients to reproduce the underlying chemical shift changes for each residue. Indeed, the data is universally well fit by this model (Fig. 2B, S9).

### Charge clustering and hydrogen bonding stabilize protonated state and shift conformational ensembles in simulations

To gain additional molecular insight into how individual ionizable residues respond to changes in pH, including transient changes in peptide conformational ensembles, we used constant pH molecular dynamics simulations of a TUBA1A^CTT^ peptide (Table S1). We performed constant pH simulations using GROMACS (58), for a total simulation time of 3 ms at pH 6 and 1 ms at pH 4.5, 5.0, 5.5, 6.5, and 7.0. We calculated the p*K*_a_ of ionizable groups in each conformation using the pypKa package (59). Using a combination of built-in and custom scripts, we identified the pairwise distance between ionizable groups, the formation and length of hydrogen bond interactions, as well as properties of the conformational ensemble, including dihedral phi/psi angles. As with the experimental data, we found that the simulation data are well fit by a two-site model, with fit pseudo p*K*_a_ values of 4.29 ± 0.03 and 5.38 ± 0.03, as compared to a p*K*_a_ of 4.25 used as the model value in the simulation for isolated glutamate residues.

Through detailed analysis of the individual trajectories, we found that both electrostatic repulsion and hydrogen bonding modulated the protonation state of glutamate side chains. The effects of electrostatic repulsion were especially apparent when we monitored the pypKa-predicted p*K*_a_ values of individual side-chain ionizable groups leading up to a protonation event. In individual examples (Fig. 2D), we observed that the predicted p*K*_a_ of two otherwise similar ionizable groups diverged as the two groups come in proximity, which results in protonation of one of the groups.

Following protonation, a hydrogen bond typically forms between the two side chains, or to a lesser extent between the protonated side chain and the backbone carbonyl. The hydrogen bond appears to stabilize the protonation, leading to a subset of protonation events that are longer than those of residues in isolation (Fig. S4). Hydrogen bond formation also stabilizes the peptide in a looped conformation, resulting in bent conformation of intervening amino acids (e.g. Fig. S6, S7).

We then sought to determine whether all ionizable side chains were equally likely to be protonated and form hydrogen bonds, or whether features of the sequence result in a preference for some amino acids. Mapping the pairwise interaction rates due to the formation of hydrogen bonds onto the sequence (Fig. 2E, S5) reveals that glutamate residues show some selectivity, with the glutamate residues in a cluster showing a higher prevalence of interactions, as expected given the proximity of other charged groups. In addition, we found that hydrogen bond interactions were most likely to be formed with non-neighboring amino acids (Fig. 2E, S5). Although pairwise hydrogen bonding interactions primarily occurs between side chain, side chain – backbone hydrogen bonding also occurs, albeit 2-3 times less often (Fig. S6).

Intra-molecular hydrogen bonding decreases the overall peptide size and thus the distance from the microtubule surface (S6). Moreover, the distribution of backbone dihedral angles is shifted in peptides with protonated side chains away from PPII, consistent with our CD measurements (Fig. 1B, S7). Taken together, our NMR, CD and simulation data clearly point to a shift in the conformational ensemble of the CTT peptide as a function of pH.

### CTT pH response is highly conserved

We then sought to determine whether the observed shift of pKa and hydrogen bonding interactions observed for the TUBA1A^CCT^ were also present in a CTT peptides taken from a range of organisms – *Homo sapiens* β/TUBB, β/TUBB3; *Caenorabditis elegans* β/MEC-7, β/TBB-4; *Saccharomyces cerevisiae* α/Tub1, β/Tub2; *Tetrahymena thermophila* α/ATU1, β/BTU1. The NMR pH titration results for CTT peptides derived from four species were similarly consistent with findings from TUBA1A^CTT^ (Table S1, Fig. 3A,B, Fig. S8-9). We found very high pH_m_ values, where roughly 99% of the variability in the chemical shifts was captured in the first two PCA components, and those components were well fit by a two-site model (Eq. 1, Fig. 3A,B, S9). We additionally fit the first PCA to a Hill equation to demonstrate the two-site character of the curves (Fig. 3B). The overall trends of values were not significantly affected by changes in buffer or magnesium concentration, nor with most minor mutations (Fig. S10). Our results indicate that the cooperative response at a higher pH than for isolated glutamates is a strongly conserved feature of tubulin CTTs, despite their underlying variability in sequence.

**Figure 3.**
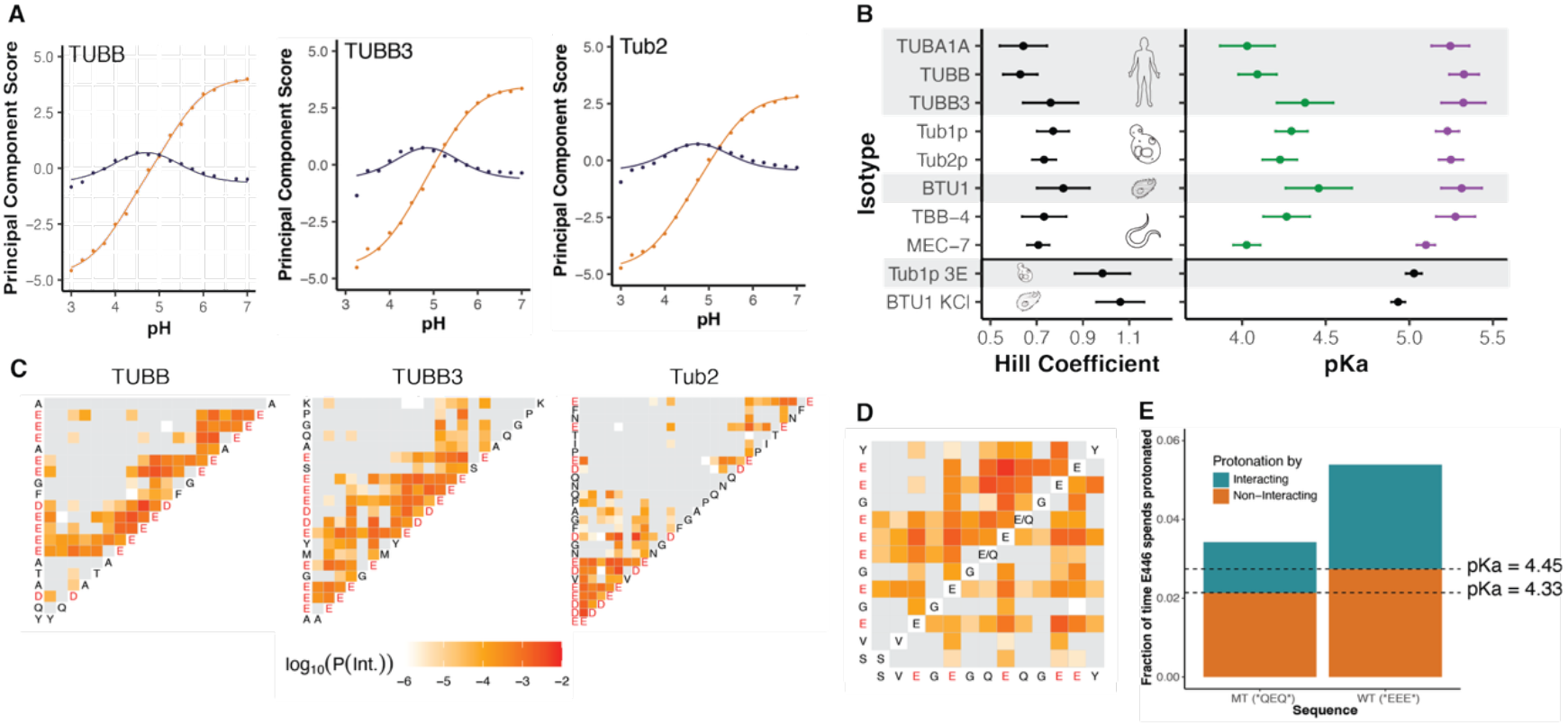
CTT pH response is conserved between species and modulated by intentional mutations. A) The first two principal components showing two dominant p*K*_a_ values. B) Fit results from either a Hill Equation or a two component Henderson-Hasselbalch equation to the first PCA of the NMR data for CTT sequences from representative eukaryotic organisms in (top) standard conditions and (bottom) with either a mutation to reduce the glutamate cluster (Tub1p-3E) or in higher salt buffer (BTU1 KCl). The experimental curves that were best fit by a two-component fit (Eq. 1), have p*K*_a1,2_ represented in green or purple respectively. Mutations to the glutamate cluster of Tub1p or addition of 150 mM KCl results in pH titration curves with reduced apparent cooperativity and better fits to a single-component Henderson-Hasselbach equation (black symbols). C) Heat maps showing interaction hotspots in glutamate clusters. D) A comparison between the wild type (top half) and QEQ mutant (bottom half) of TUBA1A^CTT^, showing the importance of neighboring charged residues on protonation. E) This effect is further quantified in a stacked bar graph showing the fraction of time E446 is protonated in QEQ and wild type sequences.

Constant pH molecular dynamics simulations of peptides taken from TUBB^CTT^, TUBB3^CTT^ and Tub2^CTT^ (Table S1) were again consistent with those of TUBA1A^CTT^, showing a higher-than-expected p*K*_a_s resulting from stabilization of protonated state by nearby charge residues and by the subsequent formation of a hydrogen bond. Thus, protonation effects resulting in altered p*K*_a_ values as observed by NMR and MD appear to be robust and universal among the isotypes and species tested.

In mapping the prevalence of hydrogen bond interactions among pairs of residues for TUBB^CTT^, TUBB3^CTT^ and Tub2^CTT^, we again found heterogeneity between pairs of interacting residues (Fig. 3C). As for TUBA1A^CCT^, we found that amino acids within glutamate clusters were more likely to form hydrogen bonds. Those residues in the middle of the largest clusters (e.g. E446 for TUBA1A, Fig. 2E or E433 for TUBB, Fig. 3B) showed the highest rates of protonation and hydrogen bond formation. As seen for TUBA1A^CCT^, the most likely partner with which these residues form a hydrogen bond was not their immediate neighbors, but residues slightly further away. We measured the frequency of hydrogen bond formation between residues as a function of residue separation and found that the most common side-chain—side-chain hydrogen bond formation was at a residue separation of four amino acids (and two amino acids for side-chain—backbone hydrogen bonds), as seen for TUBA1A^CTT^ (Fig. S6).

It appears that there are two effects at play in the pH response of CTTs. First, adjacent charged residues favor the protonation of a residue because of the electrostatic energetic cost of the high density of charged groups. This is seen by the increase in protonation rates even for non-interacting amino acids (Fig. 3E). Second, a charged residue interacts with an amino acid more distal in primary sequence. This interaction further drives protonation and hydrogen bond formation stabilizing the protonated state (Fig. 3E, S4). This may explain the relatively high prevalence of glycine residues within acidic clusters observed in CTT sequences (Fig. S11), giving chain flexibility that allows for such interactions.

### Mutations and ionic strength modulate pH response of CTTs

We sought to use these insights to modulate the behavior of the cooperative and elevated pH response. First, given that the residues showing the highest level of protonation occurred in acidic clusters of at least three amino acids (Figs. 2E, 3C), we hypothesized that isolating a residue in one of the clusters would significantly decrease the fraction of time that residue is protonated. To test this, we simulated mutant TUBA1A^CTT^ with E445Q and E447Q (TUBA1A^CTT^-QEQ). This mutation results in fewer interactions within the TUBA1A^CTT^, especially for E446 as compared to the wild type (Fig. 3D). The simulation-averaged p*K*_a_ value of E446 was 4.33 ± 0.01 in the QEQ mutant, a decrease of 0.12 pH units compared to E446 in the wild-type CTT with EEE (Fig. 3E). Moreover, the length of the protonation events is typically shorter in the mutant than the WT (Fig. S12). This reinforces the idea that the high overall charge density within polyglutamate clusters drive pairwise residue interactions for medial glutamate residues within the cluster.

We also saw a modulation of the pH response behavior in two NMR titration experiments (Fig. 3B). First, we hypothesized that we could reduce interactions by reducing the size of the glutamate cluster in the yeast alpha-tubulin CTT (Tub1^CTT^). Tub1^CCT^ only contains 5 glutamates, four of which are in a single cluster. As we had found that very few interactions occur with neighboring residues, we reasoned by reducing the size of the cluster to three that we would significantly decrease the cooperativity of the interaction. Second, we added 150 mM KCl to screen electrostatic interactions we expected to be important for the cooperativity in a titration of Tetrahymena β-tubulin, BTU1^CTT^. In both cases, the first PCA score was not well fit by a two-state model and was instead much better fit by a single site model (Fig. 3B). As further evidence for the lack of cooperativity, we fit these as well as the wild type PCA scores to a Hill equation and found fit Hill coefficients near 1 for only these two experiments among all our conditions tested. Thus, either significantly increasing the ionic strength or reducing the size of the glutamate clusters was sufficient to reduce the coupled nature of the pH response.

### CTT-tail dependent pH regulation of Cin8 binding

CTTs are known to support interactions with a variety of microtubule binding proteins. Given the high p*K*_a_ of CTT protonation and the changes in CTT conformation upon protonation, we hypothesized that CTT protonation may provide a mechanism for regulating these interactions. Previous studies by our lab and others demonstrated that the budding yeast kinesin-5 motor, Cin8, binds to the β-tubulin CTT and that this interaction promotes recruitment to microtubules and plus-end directed motility (60, 61). We used *in vitro* reconstitution and TIRF imaging to visualize the binding of 3eGFP labeled Cin8 to yeast microtubules grown from lysates of cells with either wild-type β-tubulin or with β-tubulin lacking the CTT (Fig 4)(60). While these experiments are typically done at pH 6.9, we varied the pH through the physiologically relevant range for budding yeast of 5.6 – 6.9 (11). Cin8 binds to microtubules in clusters, and so we quantified the number of clusters per unit length, the average cluster intensity, as well as the total integrated intensity per unit length (Fig. 4, Fig S13).

**Figure 4.**
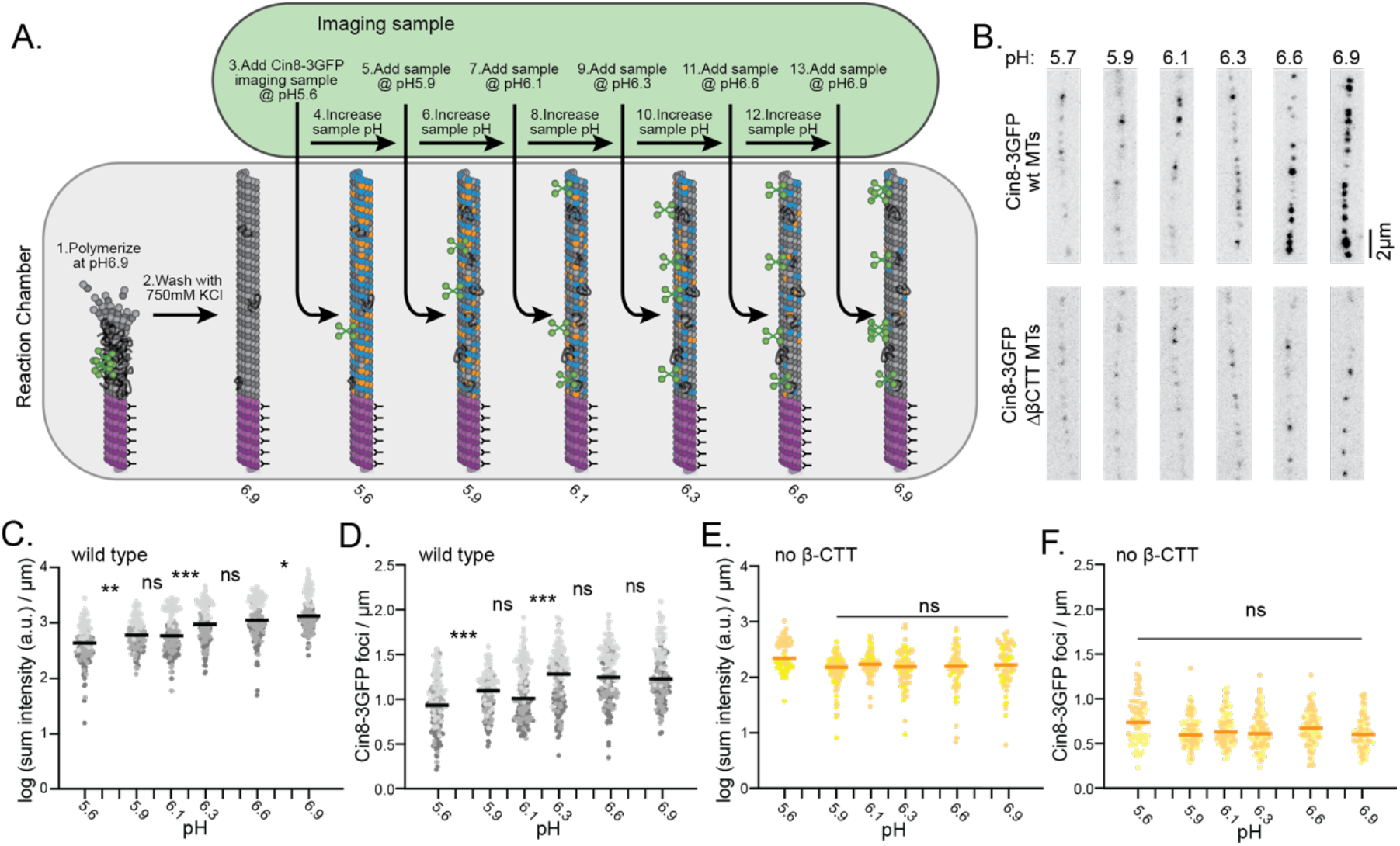
Cell lysate experiments demonstrate significant pH dependence of the binding of Cin8 to microtubes. A) An overview of the experimental approach. Dimers were randomly colored blue or orange based on the predicted fraction of monomers with at least one proton on a glutamate side chain for β- and α-tubulin respectively. B) Representative images of microtubules from each pH point. C,D) Quantification of the summed intensity along the MT length (C) and the number of foci per unit length (D) for wild-type tubulin. E,F) We repeated the same experiments for tubulin taken from cells lacking the β-tubulin CTT and saw no increase in either sum intensity or linear density of foci.

By all measures, we found that Cin8-3eGFP binding increases as pH is increased (Figure 4B-D). This increase is significantly greater than what we observed in control experiments with eGFP-labeled tubulin; we find an approximate 30% increase in eGFP-tubulin signal from pH 6 to pH 7 but a 60% increase in Cin8-3eGFP signal across the same range (Fig. 4C, S13). Therefore pH-dependent changes in eGFP fluorescence cannot explain our results for Cin8-3eGFP (62). Moreover, the number of Cin8-3eGFP foci detected along microtubules increases with pH (Fig. 4D). These results suggest that lower pH, where tubulin CTTs are more likely to be protonated, Cin8 has a lower binding affinity to microtubules.

Next, we tested whether the pH-dependent binding of Cin8-3eGFP depends on the β-tubulin CTT. We repeated the experiments using reconstituted microtubules from mutant yeast in which the β-tubulin CTT is removed by genome editing (63). Consistent with our previous results(60), we find that Cin8 binding to microtubules is decreased in the absence of β-tubulin CTT, compared to wild-type microtubule. Strikingly, the amount of Cin8-3eGFP measured on microtubules lacking β-tubulin CTT is not significantly changed across pH 5.6 – 6.9 (Figure 4E-F). These results indicate that β-tubulin CTT is necessary for the pH-sensitive binding of Cin8 to microtubules.

### CAP-Gly2 binding to TUBA1A^CTT^ shows pH-dependent chemical shift changes but not binding affinity

The CAP-Gly1 and CAP-Gly2 binding domains of CLIP170 are well characterized α-tubulin CTT binding domains. CAP-Gly binding is centered on the terminal tyrosine(64), though glutamate residues near the terminus are involved in the binding interface. Using NMR, we compared the residue-specific chemical shift perturbations of TUBA1A^CTT^ bound to the CAP-Gly domains at pH 6 and pH 7 (Fig. S14-15). We observed that the chemical shift perturbations between the two pH values are higher for the TUBA1A^CTT^ bound to CAP-Gly domains than in the apo state. This could either be due to changes in transient interactions with the CAP-Gly surface, or due to different allowable conformations in the bound and unbound states.

To determine whether these changes were sufficiently large to affect the binding affinity, we performed fluorescence anisotropy experiments of TUBA1A^CTT^ binding to the CAP-Gly2 domains. To control for any potential pH dependence of the interaction of the dye molecule itself, we measured the affinity of unlabeled TUBA1A^CTT^ by using it to compete off dye-labeled TUBA1A^CTT^ (65). Despite the difference in chemical shifts of the bound and unbound peptide in response upon changing pH, our fluorescence anisotropy experiments suggest a similar binding affinity of TUBA1A^CTT^ and CAP-Gly2 at pH 6 and 7 (Fig. S16B). Taken together, these data show that although there are differences in either the conformational ensemble or transient binding interactions of TUBA1A^CTT^, they do not affect the binding affinity between the TUBA1A^CTT^ and the CAP-Gly2 domain.

## Discussion

In this work, we used tubulin CTTs to test the hypothesis that glutamate-rich intrinsically disordered regions generally respond to changes in cellular pH within physiological pH ranges. Indeed, we found that CTTs taken from a range of organisms display a robust pH response. Across all species tested, the CTTs remain primarily disordered but show significant pH-dependent changes in conformational ensemble as measured by CD, NMR, and simulation. The experimental and simulation data are both well described by a model of two pseudo-sites, arising from interactions among protonatable residues. The resultant average p*K*_a_ values are universally elevated relative to the model p*K*_a_ values for glutamate and aspartate residues in isolation. We attribute the shift in p*K*_a_ values to the electrostatic repulsion among charged residues and the stabilization of the protonated state by hydrogen bonding. The combination of elevated p*K*_a_ values and the very high density of ionizable residues within the CTTs results in a pH response of the CTTs that is likely physiologically relevant.

Our results also suggest a molecular syntax for the enrichment of glutamate residues in tubulin CTTs that may apply to other proteins. Our modeling results indicate that pH-dependent hydrogen bonding most often occurs between glutamate residues that are separated by 3-4 residues. This suggests that pH-dependent hydrogen bonding could create a selective pressure for the retention of glutamate residue density and spacing in tubulin CTTs. Indeed, our analysis of CTTs across a wide swath of eukaryotes indicate that, in nearly all cases, at least half of the residues are glutamates and the majority feature clusters of at least 7 amino acids in length where over 70% are glutamates (Figure S17). This provides multiple pairs of glutamates separated by 3 residues, ideally positioned for hydrogen bonding in our simulations. In addition, we find that clusters of glutamate residues in CTTs often feature one or more glycine residues interspersed between glutamates (Figure S11). Glycine residues could provide the flexibility that enables intra-chain hydrogen bonds, consistent with previous work showing a decrease in pH sensitivity of titin with interspersed proline residues(66).

Previous literature has characterized tubulin CTTs as charged brushes coating the microtubule surface where they support electrostatic interactions with MAPs and alter microtubule dynamics(67–70). Our findings add to this model by suggesting that protonation alters the conformations of CTTs by creating bends through hydrogen bonding between CTT glutamate residues. We hypothesize that these bends can, very generally, create distinct conformations that could either promote or inhibit MAP binding. A bent conformation created by protonation could inhibit MAP binding by decreasing the number of glutamate side chains available for electrostatic interactions and shortening the capture radius of the CTT at the microtubule surface. This is reminiscent of our findings for the yeast kinesin-5, Cin8, which exhibits weaker binding to microtubules at low pH values where the β-tubulin CTT is more often protonated (Fig. 4). Alternatively, a bent conformation could create new binding states that support interactions with some MAPs and may be particularly relevant for MT modifying enzymes. For example, structural studies show that tubulin tails adopt a bent conformation when fit into the active site of the peptidase CCP5(71). With the plethora of known microtubule binders, protonation could act as a quick and efficient PTM for controlling a diversity of microtubule functions scaling from direct effects on MAP binding, to secondary effects created by gain or loss of those altered MAPs, and ultimately to network-scale effects on cytoskeletal architecture, trafficking, and signaling.

How might protonation compare to other CTT PTMs that also alter MAP interactions and network function? Our simulations predict that CTT protonation creates hydrogen bonding events with lifetimes on the order of 100 ps; therefore, protonation enables rapid and high frequency sampling of the modified state. In contrast, microtubule PTMs such as acetylation, glutamylation, and detyrosination/tyrosination exhibit slower dynamics, with enzyme kinetics on the order of seconds to minutes (71–74). These PTMs are controlled by specific ‘writer’ enzymes that add the modification and ‘eraser’ enzymes that remove the modification (42). Accordingly, these reactions are limited by the expression levels of the enzymes, their binding to the tubulin or microtubule substrate, and the rate of the addition or removal reaction. Despite slow dynamics, these canonical PTMs create modified states that are highly stable and accumulate on long-lived microtubules. An intriguing possibility is that CTT protonation could influence canonical PTMs by altering the activities of the modifying enzymes, or that canonical PTMs could influence protonation by modifying glutamate side chains in the CTT. We note that the glutamate residues identified in our study as either protonated or immediately adjacent to protonated sidechains are among the primary sites of polyglutamylation identified in brain tubulin (Table S1) (75–80). This creates potential for a fast signal to convert into a slow signal, and vice versa.

In conclusion, protonation can act as a quick and efficient mechanism for controlling MAPs which bind the microtubule lattice and thus regulate the diversity of functions the microtubule cytoskeleton performs, including signaling, trafficking, and cell division Given the prevalence of glutamate clusters in human proteins generally (S17), this may be a widely used mechanism of cellular pH sensing.

## Materials and Methods

As described in more detail in the SI, GST tagged C-terminal tails were expressed and purified from E. coli cells, cleaved from the tag with thrombin, and then isolated. NMR experiments were performed on an Agilent 600MHz INOVA spectrometer using standard BioPack experiments. For the NMR titration experiments, samples were started at pH 7 and then brought down to pH 3 in 0.25 pH unit increments using HCl and KOH. The added volumes of acid or base was tracked and in total did not exceed 20% of the sample volume. NMR binding experiments were performed using 15N labeled TUBA1ACTT peptide and CAP-Gly1 or CAP-Gly2.

CD experiments were done on an Applied Photophysics Chirascan Plus (Leatherhead, UK at wavelengths 188-260 (TUBB, TUBA1A) or 189.5-206 (TUBB3) and data was analyzed after buffer subtraction. Fluorescence anisotropy experiments were performed on FITC-labeled TUBA1A peptide (Genscript) using Tecan Spark plate reader. For the initial FA assay to determine the affinity of the CAP-Gly2 for FITC-TUBA1A^CTT^, fluorescently labeled TUBA1A^CTT^ was held constant while CAP-Gly2 was varied. For the unlabeled TUBA1A competition assays, FITC-TUBA1A^CTT^ and CAP-Gly2 were held constant, and unlabeled TUBA1A^CTT^ was varied(65).

The molecular dynamics experiments were performed using the GROMACS constant pH package (58, 81) using the CHARMM36 force field (82). Details of the setup and equilibration are described in the SI. Microscopy experiments were performed essentially as detailed in (60), except that the Cin8 lysate was first brought to the lowest pH, and then slowly raised in ∼0.2 pH units using KOH between subsequent experiments. Image quantification was done using MATLAB (MathWorks, Natick, MA).

## Acknowledgments

We thank Kevin Slep for plasmids encoding the CAP-Gly1/2 domains of CLIP170. We thank Emma Seidler for help with pilot NMR experiments, David Jones for help with NMR instrumentation, and Dan Sackett for useful discussions.

We thank Annette Erbse and the Shared Instruments Pool (RRID: SCR_018986) of the Department of Biochemistry at the University of Colorado Boulder for access to and support of the CD spectrometer, (NIH Shared Instrumentation Grant S10RR028036), the Avanti JXN 26 Centrifuges (NIH, R24OD033699), and the Spark Multimode Plate Reader. The Agilent 600MHz INOVA spectrometer was purchased and supported by the National Institutes of Health (NIH) (RR011969, RR16649) and the National Science Foundation (NSF) (DBI-0230966, 960241). L.H. and J.A. were supported by NSF MCB-1943488 and NIH R35GM119755, and G.F-I was supported by the University of Colorado Boulder SMART program and DOE REU PHY-2244720. J.K.M. was supported by NIH R35 GM 136253. E.C.T. was supported by the NSF Graduate Research Fellowship under Grant No.193805. P.D. and D.S. were supported by NIH R01 GM136822. We acknowledge support from NIH R01 GM136946 and R35 GM158070 to C.A.C., and postdoctoral support to N.A. from the Syracuse University Vice President of Research office. For ^13^C-detect experiments, we acknowledge support for the Bruker 800 MHz spectrometer with TCI cryoprobe at Syracuse University by shared instrumentation NIH Grant 1S10OD012254.

## Author Contributions

All authors designed research and contributed to writing and editing of the manuscript. A.W. and N.A. performed and analyzed NMR experiments, P.D. and D.S. performed and analyzed simulations, E.T., J.M and L.H. performed microscopy experiments, E.T. analyzed microscopy experiments. G.F-I and L.H. performed the bioinformatics.

## References

1. H. Hou, et al., Single-cell pH imaging and detection for pH profiling and label-free rapid identification of cancer-cells. Sci Rep 7, 1759 (2017).

2. J. R. Casey, S. Grinstein, J. Orlowski, Sensors and regulators of intracellular pH. Nat Rev Mol Cell Biol 11, 50–61 (2010).

3. L. K. Putney, D. L. Barber, Na-H Exchange-dependent Increase in Intracellular pH Times G2/M Entry and Transition *. Journal of Biological Chemistry 278, 44645–44649 (2003).

4. B. Ulmschneider, et al., Increased intracellular pH is necessary for adult epithelial and embryonic stem cell differentiation. J Cell Biol 215, 345–355 (2016).

5. X. Zhang, Y. Lin, R. J. Gillies, Tumor pH and its measurement. J Nucl Med 51, 1167–1170 (2010).

6. J. Michl, et al., Acid-adapted cancer cells alkalinize their cytoplasm by degrading the acid-loading membrane transporter anion exchanger 2, SLC4A2. Cell Reports 42 (2023).

7. J. S. Spear, K. A. White, Single-cell intracellular pH measurements reveal cell-cycle linked pH dynamics. [Preprint] (2021). Available at: https://www.biorxiv.org/content/10.1101/2021.06.04.447151v1 [Accessed 19 March 2022].

8. T. Kato, et al., Decreased brain intracellular pH measured by 31P-MRS in bipolar disorder: a confirmation in drug-free patients and correlation with white matter hyperintensity. European Archives of Psychiatry and Clinical Neurosciences 248, 301–306 (1998).

9. A. Hulikova, A. L. Harris, R. D. Vaughan-Jones, P. Swietach, Regulation of intracellular pH in cancer cell lines under normoxia and hypoxia. Journal of Cellular Physiology 228, 743–752 (2013).

10. M. C. Munder, et al., A pH-driven transition of the cytoplasm from a fluid- to a solid-like state promotes entry into dormancy. eLife Sciences 5, e09347 (2016).

11. R. P. Joyner, et al., A glucose-starvation response regulates the diffusion of macromolecules. eLife 5, e09376 (2016).

12. C. G. Triandafillou, C. D. Katanski, A. R. Dinner, D. A. Drummond, Transient intracellular acidification regulates the core transcriptional heat shock response. eLife 9, e54880 (2020).

13. R. Dechant, et al., Cytosolic pH is a second messenger for glucose and regulates the PKA pathway through V-ATPase. EMBO J 29, 2515–2526 (2010).

14. A. S. Milliken, J. H. Ciesla, S. M. Nadtochiy, P. S. Brookes, Distinct effects of intracellular vs. extracellular acidic pH on the cardiac metabolome during ischemia and reperfusion. J Mol Cell Cardiol 174, 101–114 (2023).

15. Y. Liu, K. A. White, D. L. Barber, Intracellular pH Regulates Cancer and Stem Cell Behaviors: A Protein Dynamics Perspective. Front Oncol 10, 1401 (2020).

16. A. Hulikova, A. L. Harris, R. D. Vaughan-Jones, P. Swietach, Regulation of intracellular pH in cancer cell lines under normoxia and hypoxia. Journal of Cellular Physiology 228, 743–752 (2013).

17. R. Schreiber, Ca2+ Signaling, Intracellular pH and Cell Volume in Cell Proliferation. J Membrane Biol 205, 129–137 (2005).

18. M. Tominaga, T. Tominaga, Structure and function of TRPV1. Pflugers Arch - Eur J Physiol 451, 143–150 (2005).

19. S.-P. Tsai, et al., The Effect of Protein Fusions on the Production and Mechanical Properties of Protein-Based Materials. Advanced Functional Materials 25, 1442–1450 (2015).

20. W. R. Forsyth, J. M. Antosiewicz, A. D. Robertson, Empirical relationships between protein structure and carboxyl pKa values in proteins. Proteins: Structure, Function, and Bioinformatics 48, 388–403 (2002).

21. C. A. Castañeda, et al., Molecular determinants of the pKa values of Asp and Glu residues in staphylococcal nuclease. Proteins: Structure, Function, and Bioinformatics 77, 570–588 (2009).

22. A. Nicolli, V. Petronilli, P. Bernardi, Modulation of the mitochondrial cyclosporin Asensitive permeability transition pore by matrix pH. Evidence that the pore open-closed probability is regulated by reversible histidine protonation. Biochemistry 32, 4461–4465 (1993).

23. P. A. Bullough, F. M. Hughson, J. J. Skehel, D. C. Wiley, Structure of influenza haemagglutinin at the pH of membrane fusion. Nature 371, 37–43 (1994).

24. N. Sriwilaijaroen, Y. Suzuki, Molecular basis of the structure and function of H1 hemagglutinin of influenza virus. Proceedings of the Japan Academy, Series B 88, 226–249 (2012).

25. J. Srivastava, et al., Structural model and functional significance of pH-dependent talin– actin binding for focal adhesion remodeling. Proceedings of the National Academy of Sciences 105, 14436–14441 (2008).

26. J. J. Bravo-Cordero, M. A. O. Magalhaes, R. J. Eddy, L. Hodgson, J. Condeelis, Functions of cofilin in cell locomotion and invasion. Nat Rev Mol Cell Biol 14, 405–415 (2013).

27. A. Schönichen, B. A. Webb, M. P. Jacobson, D. L. Barber, Considering Protonation as a Posttranslational Modification Regulating Protein Structure and Function. Annual Review of Biophysics 42, 289–314 (2013).

28. K. A. White, B. K. Grillo-Hill, D. L. Barber, Cancer cell behaviors mediated by dysregulated pH dynamics at a glance. J Cell Sci 130, 663–669 (2017).

29. J. A. Read, V. J. Winter, C. M. Eszes, R. B. Sessions, R. L. Brady, Structural basis for altered activity of M- and H-isozyme forms of human lactate dehydrogenase. Proteins 43, 175–185 (2001).

30. K. P. Kisor, D. G. Ruiz, M. P. Jacobson, D. L. Barber, A role for pH dynamics regulating transcription factor DNA-binding selectivity. Nucleic Acids Res 53, gkaf474 (2025).

31. C. Valéry, et al., Atomic view of the histidine environment stabilizing higher-pH conformations of pH-dependent proteins. Nat Commun 6, 7771 (2015).

32. R. Calinsky, Y. Levy, A pH-Dependent Coarse-Grained Model for Disordered Proteins: Histidine Interactions Modulate Conformational Ensembles. J. Phys. Chem. Lett. 15, 9419–9430 (2024).

33. B. Gabryelczyk, et al., Hydrogen bond guidance and aromatic stacking drive liquid-liquid phase separation of intrinsically disordered histidine-rich peptides. Nat Commun 10, 5465 (2019).

34. M. R. King, et al., Macromolecular condensation organizes nucleolar sub-phases to set up a pH gradient. Cell 187, 1889-1906.e24 (2024).

35. J. Guillén-Boixet, et al., RNA-Induced Conformational Switching and Clustering of G3BP Drive Stress Granule Assembly by Condensation. Cell 181, 346-361.e17 (2020).

36. T. M. Franzmann, et al., Phase separation of a yeast prion protein promotes cellular fitness. Science 359, eaao5654 (2018).

37. L. Baidya, G. Reddy, pH Induced Switch in the Conformational Ensemble of Intrinsically Disordered Protein Prothymosin-α and Its Implications for Amyloid Fibril Formation. J. Phys. Chem. Lett. 13, 9589–9598 (2022).

38. V. N. Uversky, et al., Natively Unfolded Human Prothymosin α Adopts Partially Folded Collapsed Conformation at Acidic pH. Biochemistry 38, 15009–15016 (1999).

39. K. M. Ruff, et al., Molecular grammars of predicted intrinsically disordered regions that span the human proteome. Cell S0092-8674(25)01191–2 (2025). 10.1016/j.cell.2025.10.019.

40. T. K. Rostovtseva, et al., Tubulin binding blocks mitochondrial voltage-dependent anion channel and regulates respiration. Proceedings of the National Academy of Sciences 105, 18746–18751 (2008).

41. A. Schönichen, B. A. Webb, M. P. Jacobson, D. L. Barber, Considering Protonation as a Posttranslational Modification Regulating Protein Structure and Function. Annual Review of Biophysics 42, 289–314 (2013).

42. C. Janke, The tubulin code: Molecular components, readout mechanisms, and functions. J Cell Biol 206, 461–472 (2014).

43. K. M. Ruff, et al., Molecular grammars of intrinsically disordered regions that span the human proteome. [Preprint] (2025). Available at: http://biorxiv.org/lookup/doi/10.1101/2025.02.27.640591 [Accessed 23 July 2025].

44. J. L. S. Lopes, A. J. Miles, L. Whitmore, B. A. Wallace, Distinct circular dichroism spectroscopic signatures of polyproline II and unordered secondary structures: applications in secondary structure analyses. Protein Sci 23, 1765–1772 (2014).

45. M. E. Tomasso, M. J. Tarver, D. Devarajan, S. T. Whitten, Hydrodynamic Radii of Intrinsically Disordered Proteins Determined from Experimental Polyproline II Propensities. PLOS Computational Biology 12, e1004686 (2016).

46. G. Nagy, M. Igaev, N. C. Jones, S. V. Hoffmann, H. Grubmüller, SESCA: Predicting Circular Dichroism Spectra from Protein Molecular Structures. J. Chem. Theory Comput. 15, 5087–5102 (2019).

47. M. J. Fossat, R. V. Pappu, q-Canonical Monte Carlo Sampling for Modeling the Linkage between Charge Regulation and Conformational Equilibria of Peptides. J. Phys. Chem. B 123, 6952–6967 (2019).

48. M. J. Fossat, A. E. Posey, R. V. Pappu, Uncovering the Contributions of Charge Regulation to the Stability of Single Alpha Helices. ChemPhysChem 24, e202200746 (2023).

49. J. Santos, V. Iglesias, C. Pintado, J. Santos-Suárez, S. Ventura, DispHred: A Server to Predict pH-Dependent Order–Disorder Transitions in Intrinsically Disordered Proteins. International Journal of Molecular Sciences 21, 5814 (2020).

50. E. Nogales, S. Grayer Wolf, I. A. Khan, R. F. Ludueña, K. H. Downing, Structure of tubulin at 6.5 Å and location of the taxol-binding site. Nature 375, 424–427 (1995).

51. K. P. Wall, et al., Molecular Determinants of Tubulin’s C-Terminal Tail Conformational Ensemble. ACS Chem Biol 11, 2981–2990 (2016).

52. H. Okazaki, et al., Using 1HN amide temperature coefficients to define intrinsically disordered regions: An alternative NMR method. Protein Science 27, 1821–1830 (2018).

53. A. Bundi, K. Wüthrich, Use of amide 1H-nmr titration shifts for studies of polypeptide conformation. Biopolymers 18, 299–311 (1979).

54. C. J. Craven, et al., Complexes Formed between Calmodulin and the Antagonists J-8 and TFP in Solution. Biochemistry 35, 10287–10299 (1996).

55. K. Sakurai, Y. Goto, Principal component analysis of the pH-dependent conformational transitions of bovine β-lactoglobulin monitored by heteronuclear NMR. Proceedings of the National Academy of Sciences 104, 15346–15351 (2007).

56. A. Onufriev, D. A. Case, G. M. Ullmann, A Novel View of pH Titration in Biomolecules. Biochemistry 40, 3413–3419 (2001).

57. A. R. Klingen, E. Bombarda, G. M. Ullmann, Theoretical investigation of the behavior of titratable groups in proteins. Photochem. Photobiol. Sci. 5, 588–596 (2006).

58. N. Aho, et al., Scalable Constant pH Molecular Dynamics in GROMACS. J. Chem. Theory Comput. 18, 6148–6160 (2022).

59. P. B. P. S. Reis, D. Vila-Viçosa, W. Rocchia, M. Machuqueiro, PypKa: A Flexible Python Module for Poisson–Boltzmann-Based pKa Calculations. J. Chem. Inf. Model. 60, 4442–4448 (2020).

60. E. C. Thomas, J. K. Moore, Selective regulation of kinesin-5 function by β-tubulin carboxy-terminal tails. Journal of Cell Biology 224, e202405115 (2024).

61. S. K. Singh, et al., Noncanonical interaction with microtubules via the N-terminal nonmotor domain is critical for the functions of a bidirectional kinesin. Sci Adv 10, eadi1367 (2024).

62. G. H. Patterson, S. M. Knobel, W. D. Sharif, S. R. Kain, D. W. Piston, Use of the green fluorescent protein and its mutants in quantitative fluorescence microscopy. Biophysical Journal 73, 2782–2790 (1997).

63. J. Aiken, et al., Genome-wide analysis reveals novel and discrete functions for tubulin carboxy-terminal tails. Curr Biol 24, 1295–1303 (2014).

64. M. Mishima, et al., Structural basis for tubulin recognition by cytoplasmic linker protein 170 and its autoinhibition. Proceedings of the National Academy of Sciences 104, 10346–10351 (2007).

65. E. T. Usher, et al., Intrinsically disordered substrates dictate SPOP subnuclear localization and ubiquitination activity. Journal of Biological Chemistry 296, 100693 (2021).

66. S. Manukian, G. E. Lindberg, E. Punch, S. P. D. Mudiyanselage, M. J. Gage, pH-Dependent Compaction of the Intrinsically Disordered Poly-E Motif in Titin. Biology 11, 1302 (2022).

67. N. Tajielyato, L. Li, Y. Peng, J. Alper, E. Alexov, E-hooks provide guidance and a soft landing for the microtubule binding domain of dynein. Sci Rep 8, 13266 (2018).

68. K. L. Sheldon, P. A. Gurnev, S. M. Bezrukov, D. L. Sackett, Tubulin tail sequences and post-translational modifications regulate closure of mitochondrial voltage-dependent anion channel (VDAC). J Biol Chem 290, 26784–26789 (2015).

69. J. Chen, et al., α-tubulin tail modifications regulate microtubule stability through selective effector recruitment, not changes in intrinsic polymer dynamics. Dev Cell 56, 2016-2028.e4 (2021).

70. A. Roll-Mecak, Intrinsically disordered tubulin tails: complex tuners of microtubule functions? Semin Cell Dev Biol 37, 11–19 (2015).

71. J. Chen, et al., Tubulin code eraser CCP5 binds branch glutamates by substrate deformation. Nature 631, 905–912 (2024).

72. C. Egoldt, M.-C. Velluz, J. Tran, C. Aumeier, Microtubule lattice conformation and integrity regulate α-tubulin acetylation. [Preprint] (2025). Available at: https://www.biorxiv.org/content/10.1101/2025.09.09.675099v1 [Accessed 16 February 2026].

73. Y. Yue, T. Hotta, R. Ohi, K. J. Verhey, MATCAP1 preferentially binds an expanded tubulin conformation to generate detyrosinated and ΔC2 α-tubulin. bioRxiv 2025.08.14.670257 (2025). 10.1101/2025.08.14.670257.

74. Y. Yue, T. Hotta, T. Higaki, K. J. Verhey, R. Ohi, Microtubule detyrosination by VASH1/SVBP is regulated by the conformational state of tubulin in the lattice. Curr Biol 33, 4111-4123.e7 (2023).

75. V. Redeker, J. Rossier, A. Frankfurter, Posttranslational modifications of the C-terminus of alpha-tubulin in adult rat brain: alpha 4 is glutamylated at two residues. Biochemistry 37, 14838–14844 (1998).

76. J. Mary, V. Redeker, J. P. Le Caer, J. C. Promé, J. Rossier, Class I and IVa beta-tubulin isotypes expressed in adult mouse brain are glutamylated. FEBS Lett 353, 89–94 (1994).

77. M. Rüdiger, U. Plessman, K. D. Klöppel, J. Wehland, K. Weber, Class II tubulin, the major brain beta tubulin isotype is polyglutamylated on glutamic acid residue 435. FEBS Lett 308, 101–105 (1992).

78. V. Redeker, R. Melki, D. Promé, J. P. Le Caer, J. Rossier, Structure of tubulin C-terminal domain obtained by subtilisin treatment. The major alpha and beta tubulin isotypes from pig brain are glutamylated. FEBS Lett 313, 185–192 (1992).

79. B. Eddé, et al., Posttranslational glutamylation of alpha-tubulin. Science 247, 83–85 (1990).

80. J. E. Alexander, et al., Characterization of posttranslational modifications in neuron-specific class III beta-tubulin by mass spectrometry. Proc Natl Acad Sci U S A 88, 4685–4689 (1991).

81. M. J. Abraham, et al., GROMACS: High performance molecular simulations through multi-level parallelism from laptops to supercomputers. SoftwareX 1–2, 19–25 (2015).

82. R. B. Best, et al., Optimization of the Additive CHARMM All-Atom Protein Force Field Targeting Improved Sampling of the Backbone ϕ, ψ and Side-Chain χ1 and χ2 Dihedral Angles. J. Chem. Theory Comput. 8, 3257–3273 (2012).

