## Supplemental Material for "Tubulin C-terminal tails are pH sensors that regulate microtubule function"

**This PDF file includes:**

Figures S1 to S17
Tables S1
Supporting text
SI References

**Other supporting materials for this manuscript include the following:**

NMR data will be deposited at https://scholar.colorado.edu/

**Supporting Figures**

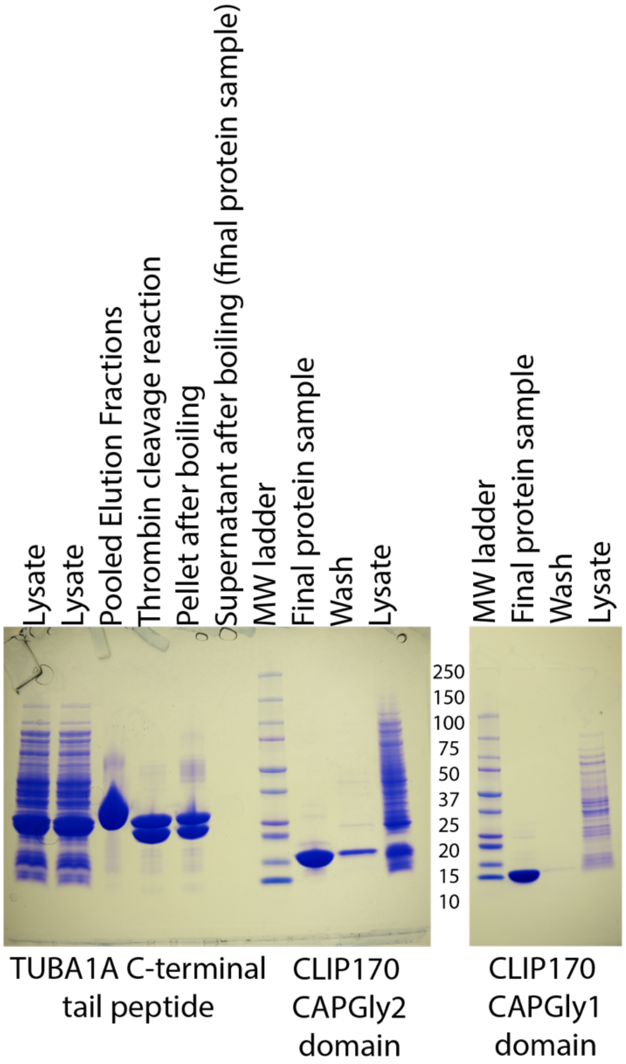


**Fig. S1.** Coomassie-stained SDS-PAGE gels of the purifications of the TUBA1A C-terminal tail peptide, and the CAP-Gly1 and CAP-Gly2 domains of CLIP170. The CTT peptides themselves are not visible on the gel due to small size and the poor binding of their amino acids to protein stains.

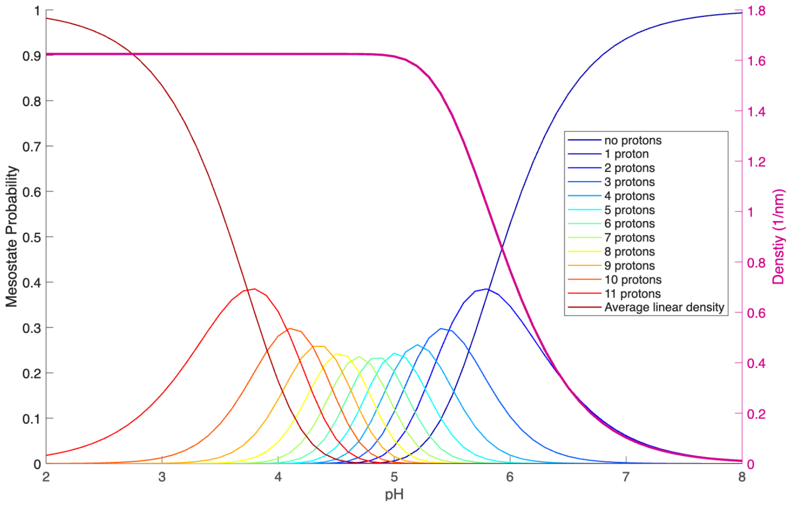


**Fig. S2.** (left axis) Mesostate probability distribution of charge states of a peptide with 11 ionizable residues, each with a p$K_a$ of 4.78. (right axis) A microtubule with 13 protofilaments and a spacing of 8 nm per tubulin dimer will have an average linear density of peptides with at least one protonated glutamate along the protofilament axis of d = (13*(1-$p_{zero}$)/8 nm), where $p_{zero}$ is the probability of having no protonated glutamates. At pH of 6.5, relevant for cancer and ischemia, the linear density is approximately 0.3 protonated glutamates/nm, or one charged glutamate per 3.3 nm. Note the spacing between dimers along a single protofilament is 8 nm.

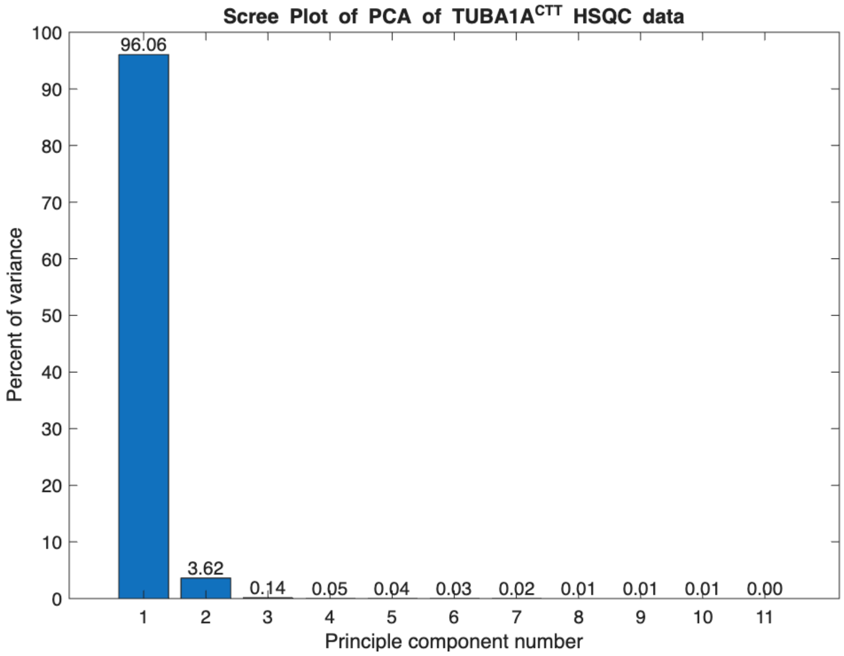


**Fig. S3.** PCA scree plot of the TUBA1ACTT chemical shift values as a function of pH. Nitrogen and proton chemical shifts were combined into a single array for analysis.

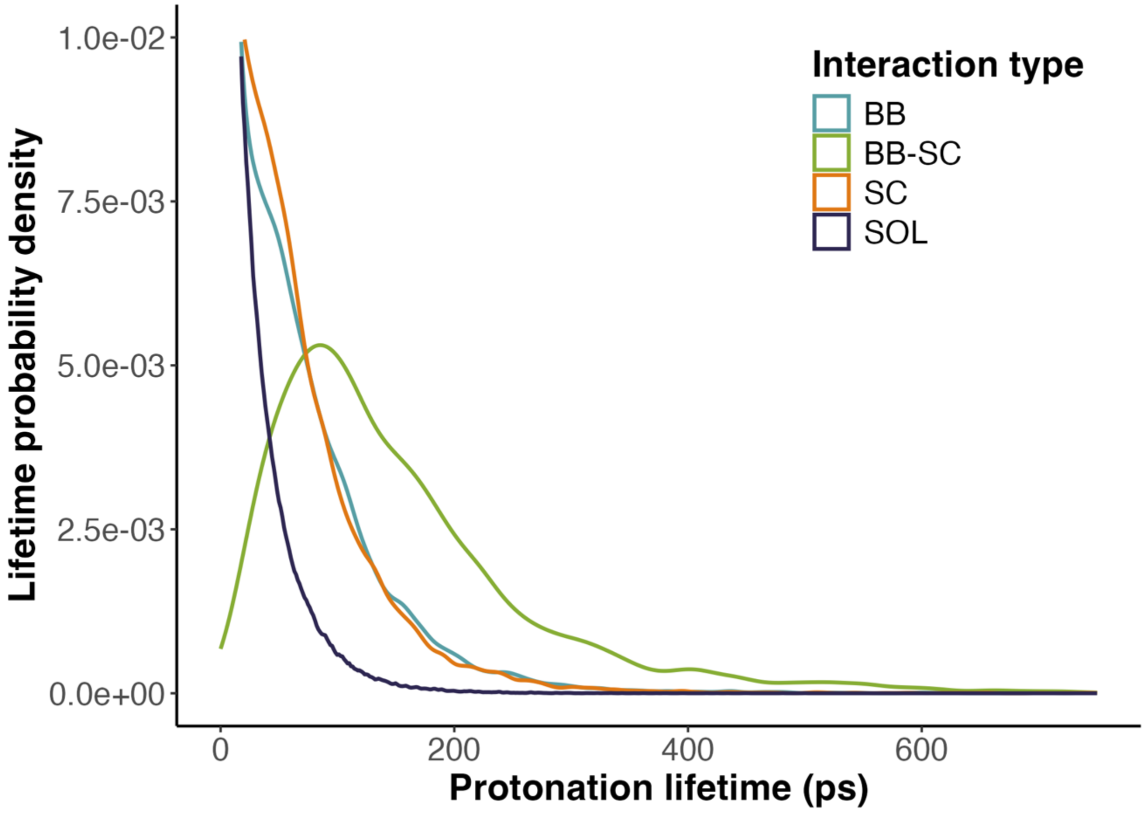


**Fig. S4.** Lifetimes of hydrogen bonding events in TUBA1A at all pHs sampled from simulations. Interactions are colored by side-chain—backbone (blue, BB), side-chain—side-chain (orange, SC), stochastic solvent protonation (purple, SOL), and combination of side-chain—backbone and side-chain—side-chain (green, BB-SC), which is where a residue will remain protonated and sequentially switch between being hydrogen bonded to a side-chain and a backbone oxygen, or vice versa.

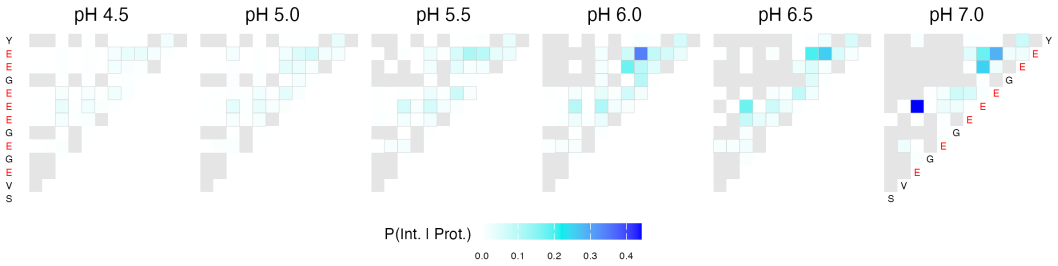


**Fig. S5.** Heatmap of pairwise residue interactions in TUBA1A at varying pH. TUBA1A sequence is displayed along the y-axis and the diagonal of the heatmaps, with acidic residues in red. Squares representing non-interacting pairs are in grey, and color bar represents the conditional probability of two residues interacting given the condition that one is protonated. This is analogous to a deviation from the Hendersen-Hasselbach (HH) equation, with a value of P(Int.|Prot.) of 0 being in line with HH prediction.

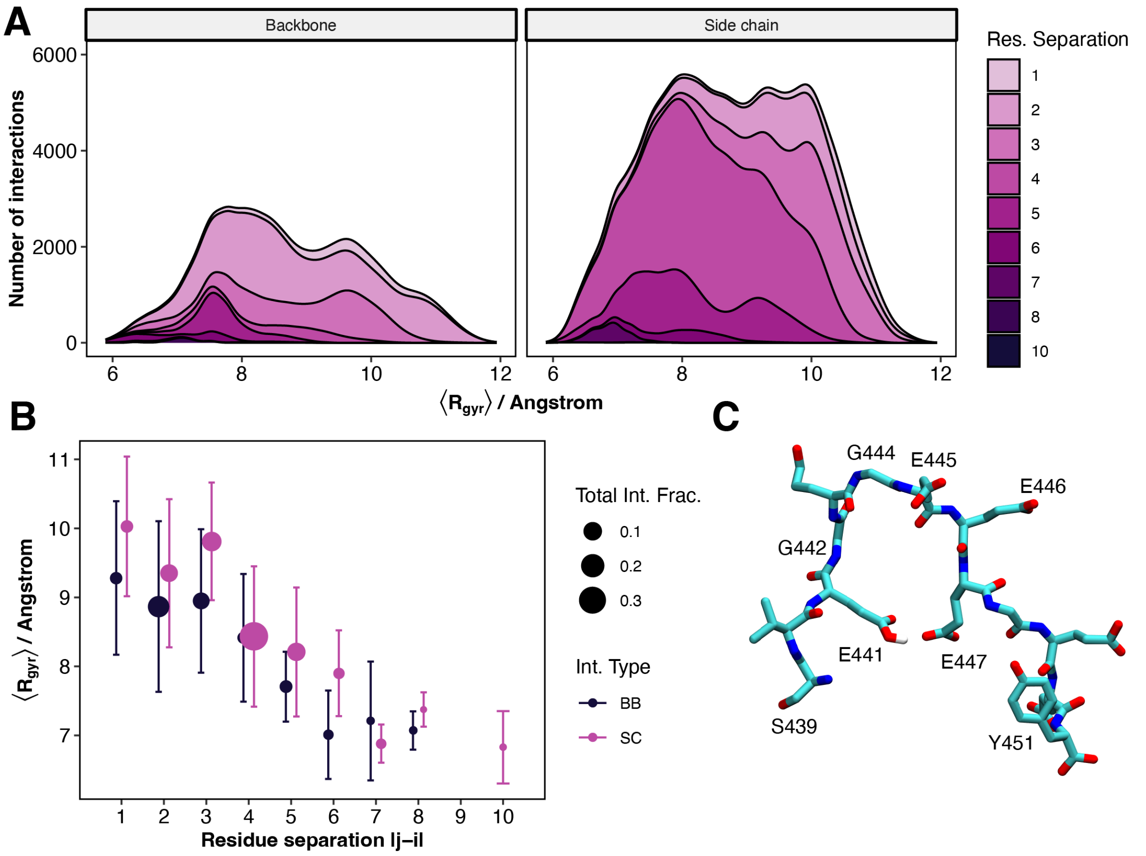


**Fig. S6.** (A) Stacked density distributions of TUBA1A$^{CTT}$ radius of gyration during hydrogen bonding events where the protonated residue forms a hydrogen bond with a backbone or side chain atom. Color represents the sequence position separation of the two residues. (B) TUBA1A$^{CTT}$ radius of gyration as a function of residue separation. Point size represents the fraction of total interactions. Hydrogen bonding events between two residue side chains four positions apart represent the largest fraction of interactions, as indicated by the largest area in A(right) and B. (C) TUBA1A$^{CTT}$ conformation taken from MD simulation showing a hydrogen bond formed between E441 and E447, a residue separation of six.

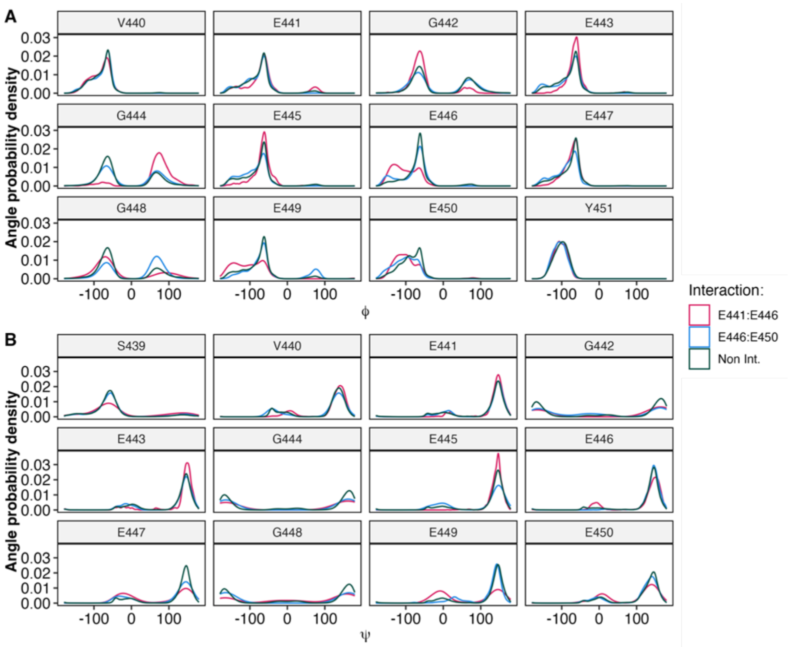


**Fig. S7.** Angle probability densities of backbone dihedral angles phi (A) and psi (B) per residue in TUBA1A during interacting (red, blue) and non-interacting (green) snapshots from MD simulations. We observe shifts in angle densities of intermediate residues between interacting pairs compared to non-interacting densities.

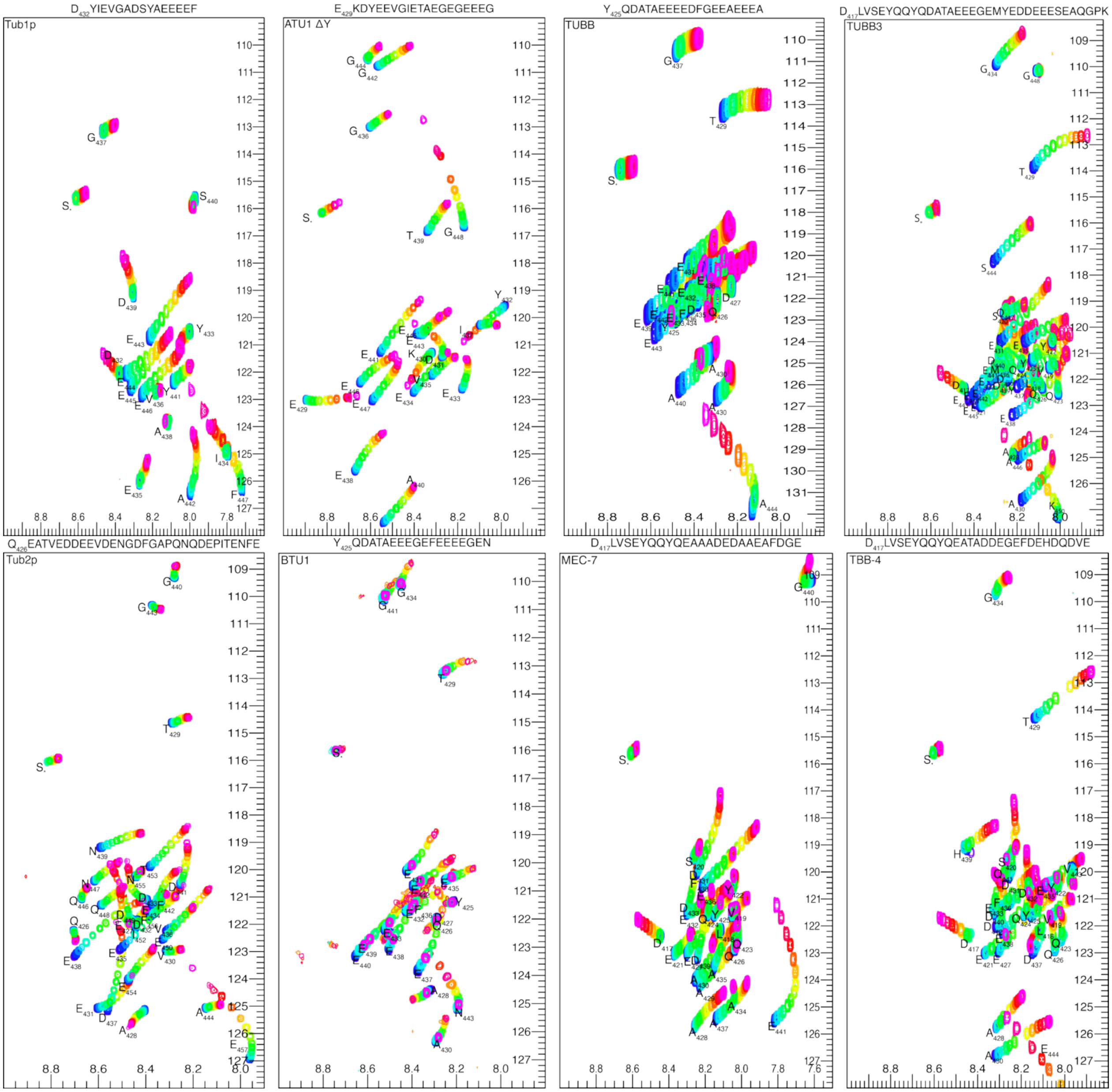


**Fig. S8.** NMR Titration experiments of CTT peptides. The protein name is listed in the top left, and the unique portion of the peptide is listed at the top, with the appropriate numbering for each organism. Each peptide begins with a GS protease scar, and the peak associated with the serine is typically visible at low pH ($S_0$). The coloring is the same as in Fig. 1. Residues with significant curvature are typically near the C-terminus of the protein. Those with more minor curvature are typically near both E and D residues.

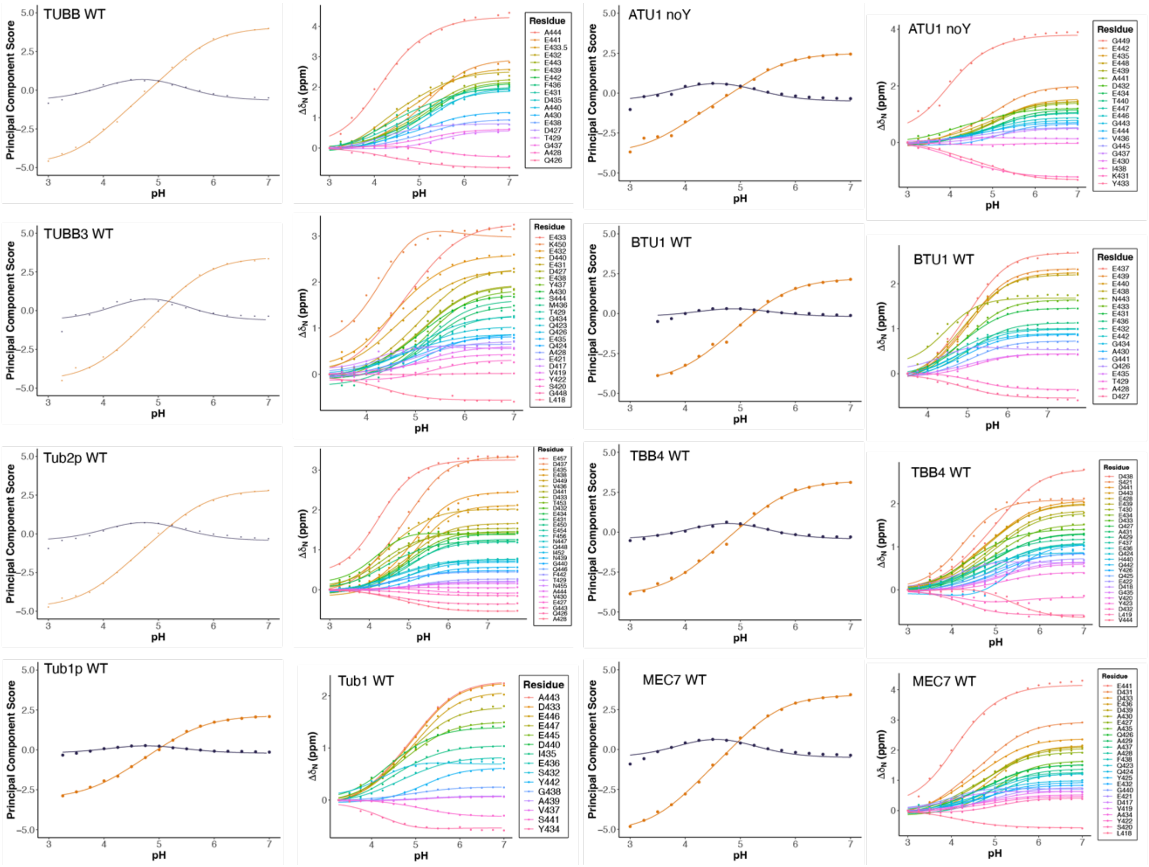


**Fig. S9.** PCA fits to the NMR chemical shift values all the different peptides showing both the fits to the first two principal components (left) and the subsequent high-quality fits to the individual chemical shifts (right) using the PCA fits for each amino acid for the peptides in Fig S8.

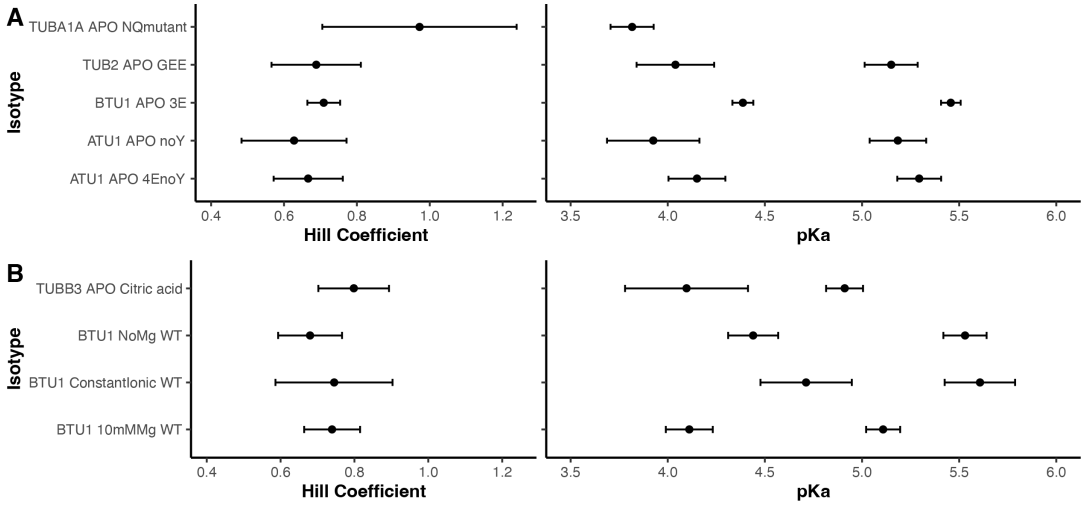


**Fig. S10.** Hill coefficient and p$K_a$ fits to the NMR chemical shift data of peptides with a variety of different (A) mutations and (B) buffers, showing that the peptide behavior is robust. Errors bars are uncertainties of the fit. The N/Q mutant has all D/E amino acids replaced with N/Q, and so the only pH responsive element is the C-terminus.

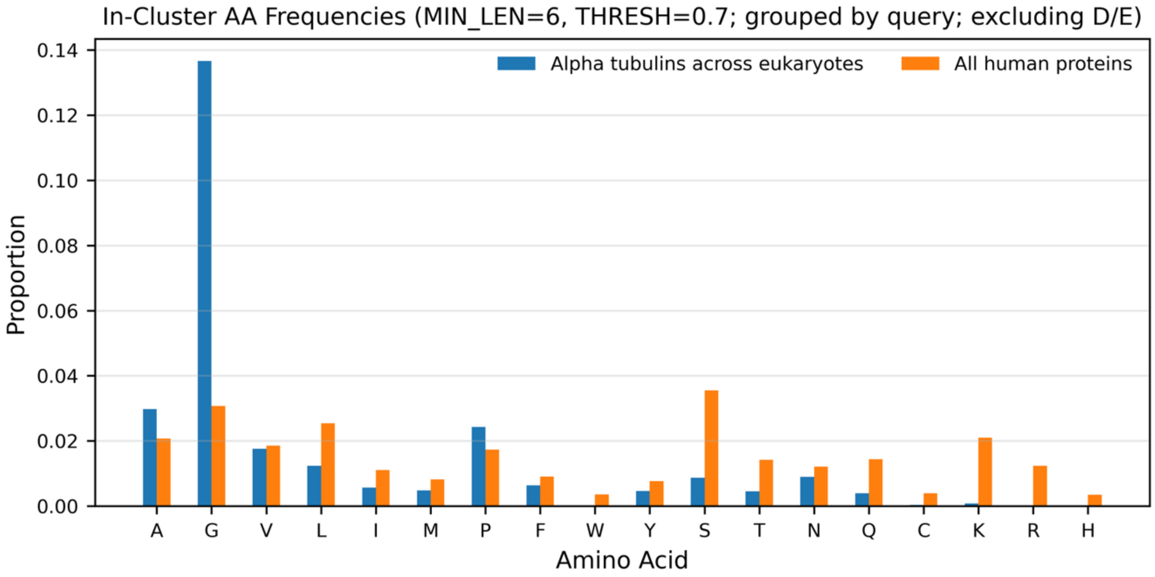


**Fig. S11**. Histogram of non-acidic residues in amino acid clusters of at least 6 amino acids in length and at least 70% acidic residues, showing a much higher prevalence of glycine residues in α-tubulin tails across eukaryotes (1) as compared to the human proteome.

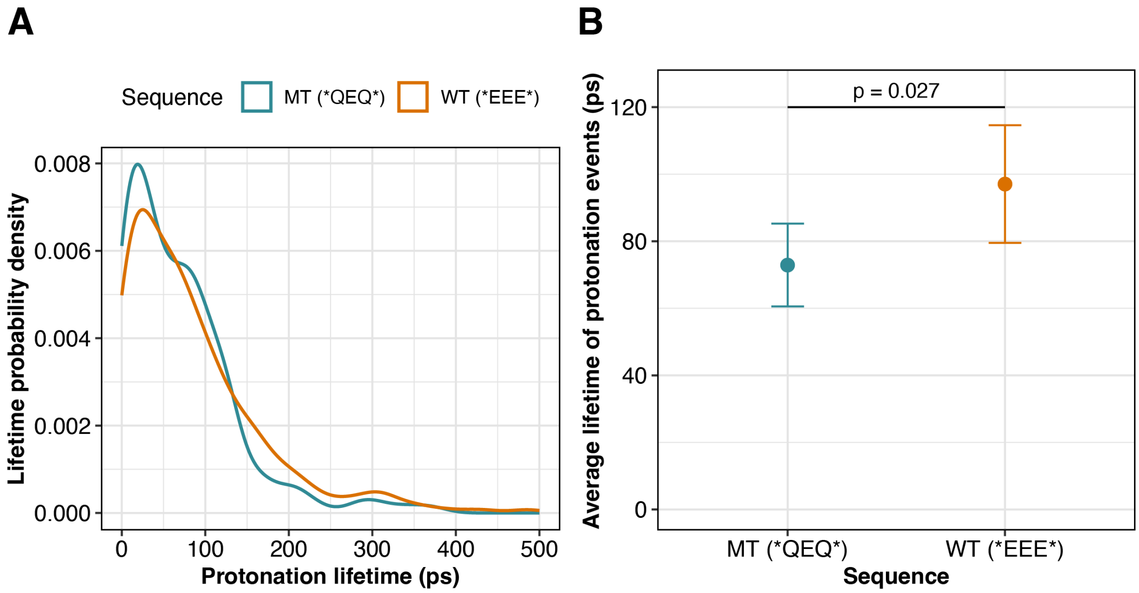


**Fig. S12.** (A) Probability density of protonation lifetimes for the WT and QEQ mutant. (B) The average lifetime of events showing that those for the wild type (EEE) are significant longer than for the mutant (QEQ).

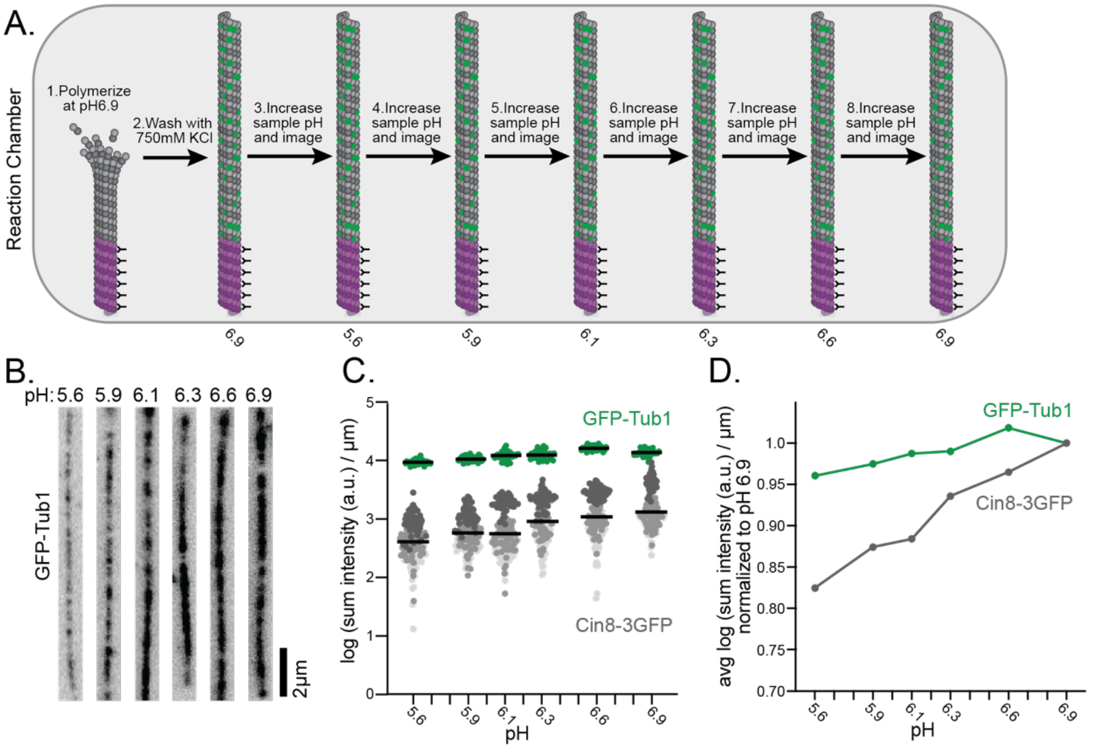


**Fig. S13.** Schematic and results from experiments with 3xeGFP labeled tubulin to show that the changes in Cin8-3eGFP are larger than that for eGFP itself. A) Schematic of the experimental setup. B) Images of microtubules at different pH (note these are different microtubules). C) Quantification of the sum intensity along the microtubule at different pH values, the Cin8-3GFP data is the same as in Fig 4. D) The same data as in Figure C normalized to pH 6.9.

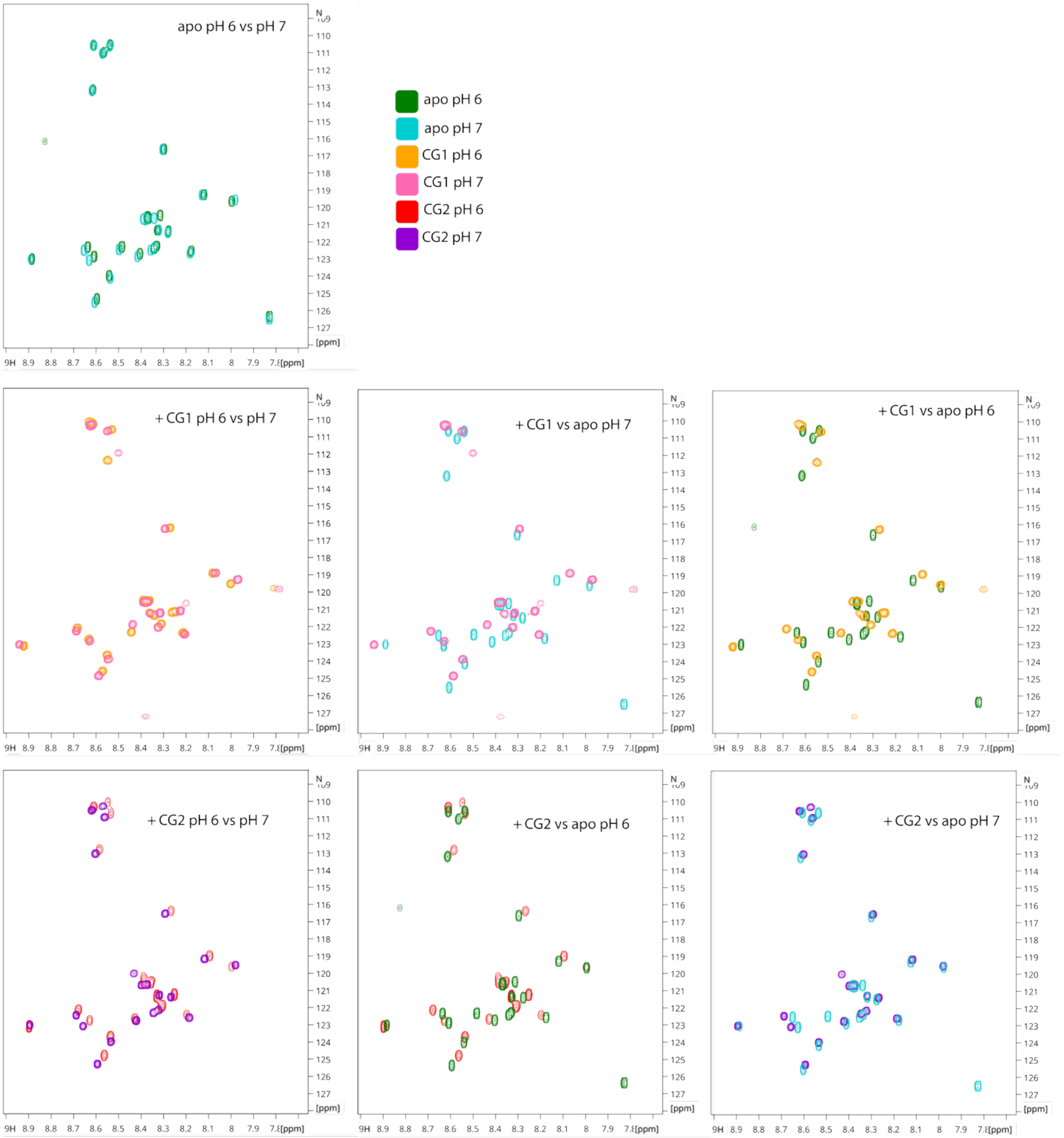


**Fig. S14.** TUBA1A$^{CTT}$ spectra with and without CAP-Gly1/2 at pH 6 or 7 as indicated showing larger changes between pH 6 and 7 in the presence of the CAP-Gly1/2 domains of CLIP170.

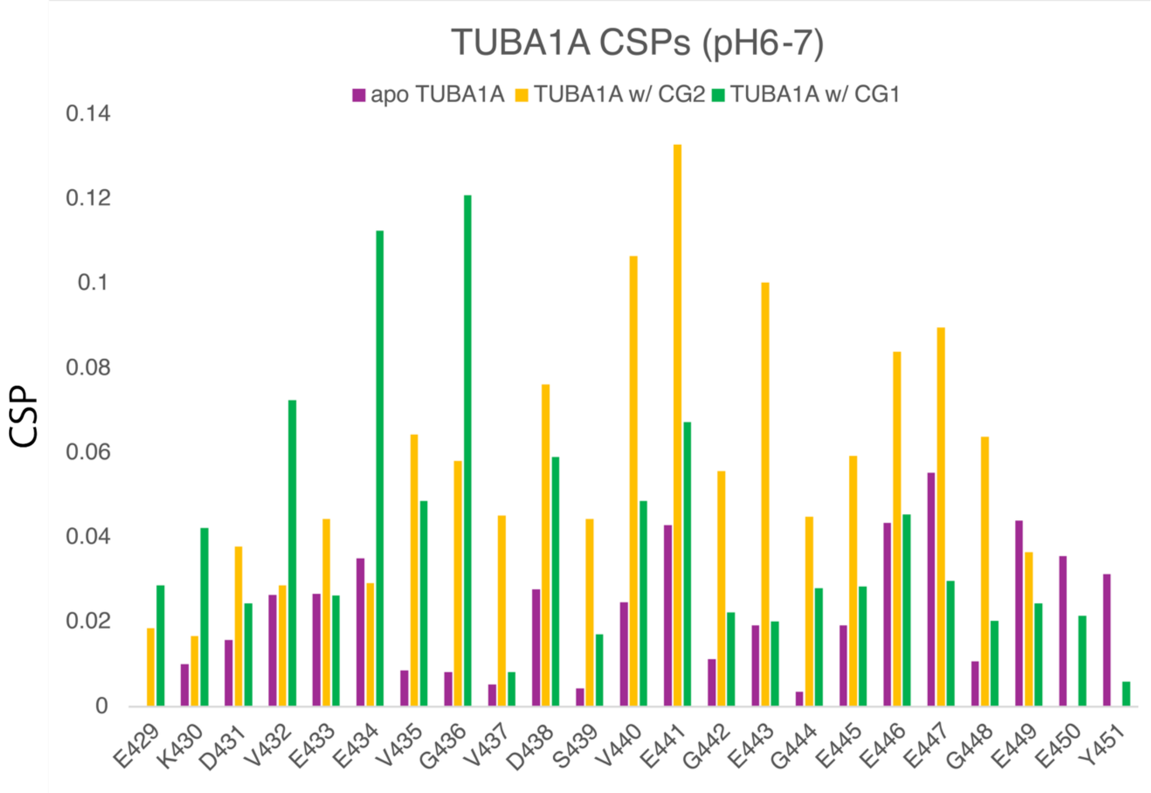


**Fig. S15.** TUBA1ACTT chemical shift perturbations for each residue between pH 6 and 7 in three conditions: APO (purple), with CAP-Gly2 domain (yellow) or with the CAP-Gly1 domain of CLIP170 (green)**.** Note that the assignment experiments were not repeated in the bound state; assignments were transferred from the APO condition.

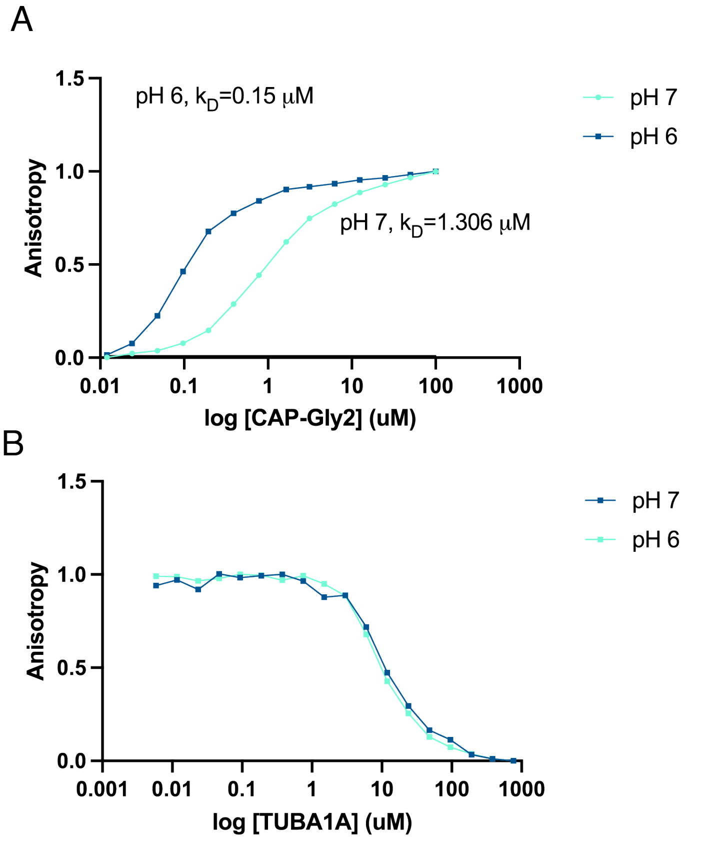


**Fig. S16.** Fluorescence anisotropy data for the binding of TUBA1A[CTT] to CAP-Gly2 (156-279) from CLIP170. The upper figure is for the labeled TUBA1A binding showing an apparent difference in $K_D$ as a function of pH. However, the pH dependence disappears in the competition assay using unlabeled TUBA1A which show no change in $K_D$ at pH 7 and 6.

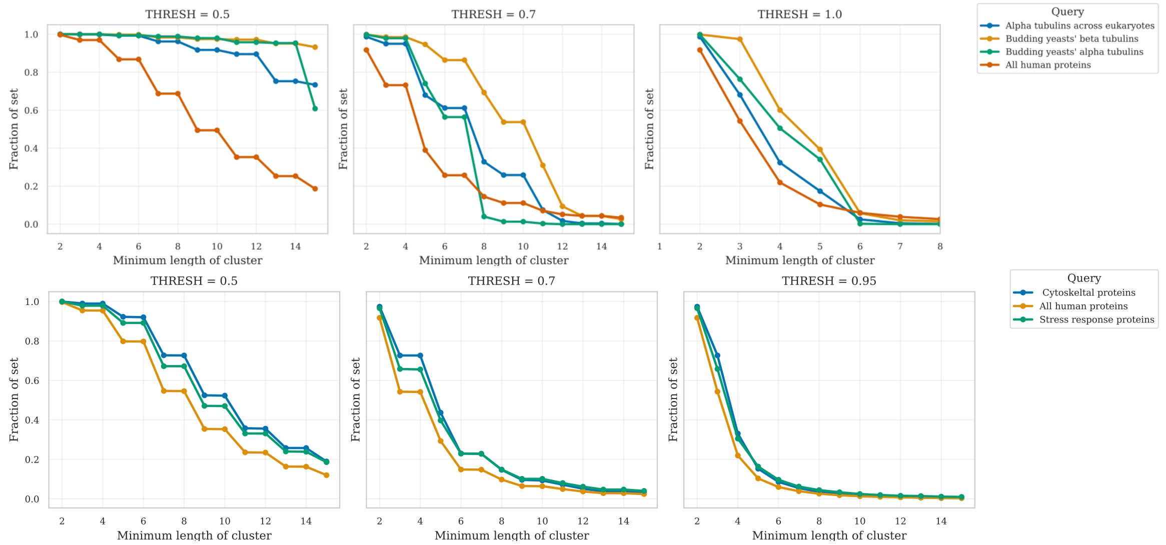


**Fig. S17.** Prevalence of acidic clusters in different groups of homologous proteins, or of human proteins with different annotations in UniProt. The different datasets are 1503 α-tubulin CTTs from across eukaryotes(1), orthogroups 775 and 651 from the yeast 1000 genomes project (2), and queries to UniProt for the human proteome (UP000005640&reviewed:true) and with the listed keywords.

| | Protein | Organism | Sequence | Amino acids | PI |
|---|---|---|---|---|---|
| **α-tubulins** | TUBA1A | H. sapiens | $E_{429}$KDYEEVGVD[SVEGEGEEEGEEY] | 23 [13] | 3.52 |
| | Tub1p | S. cerevisiae | $D_{432}$YIEVGADSYAEEEEF | 16 | 3.3 |
| | ATU1 | T. thermophila | $E_{429}$KDYEEVGIETAEGEGEEEG | 20 | 3.63 |
| **β-tubulins** | TUBB | H. sapiens | [$Y_{425}$QDATAEEEEDFGEEAEEEA] | 20 [20] | 3.18 |
| | TUBB3 | H. sapiens | $D_{417}$LVSEYQQYQDAT[AEEEGEMYEDDEEESEAQGPK] | 34 [21] | 3.41 |
| | Tub2p | S. cerevisiae | $Q_{426}$EATV[EDDEEVDENGDFGAPQNQDEPITENFE] | 32 [27] | 3.05 |
| | BTU1 | T. thermophila | $Y_{425}$QDATAEEEGEFEEEEGEN | 19 | 3.25 |
| | MEC-7 | C. elegans | $D_{417}$LVSEYQQYQEAAADEDAAEAFDGE | 25 | 3.17 |
| | TBB-4 | C. elegans | $D_{417}$LVSEYQQYQEATADDEGEFDEHDQDVE | 28 | 3.4 |

**Table S1.** List of peptides used in this study. All were produced as GST-tagged proteins and cleaved using thrombin. Each peptide begins with a GS scar from the cleavage reaction. The peptide sequence used for experimental studies is shown, with the sequence used for the molecular dynamics simulations indicated with brackets. Those residues highlighted in red are sites of PTM in brain tubulin (3–8).

**SI Extended Methods**:

CTT expression and purification

All CTT constructs were expressed heterologously in *E. coli* BL21 DE3 cells using pGEXT 4T-3 which includes a GST purification tag. Unlabeled cultures were grown to OD 0.6-0.8 in LB and induced at 23C for 16 hours with 1 mM IPTG. Labeled cultures for NMR were grown in LB to OD 0.6-0.8, washed in M9 salts (6.2 g/L $Na_2HPO_4$, 3 g/L $KH_2PO_4$, 0.5 g/L NaCl pH 7.4), resuspended in M9 media containing $^{15}NH_4Cl$ and/or $^{13}C$-glucose, and induced at 23C for 16 hours with 1 mM IPTG. Purifications were typically done on 1 L of cells, though higher for some double labeled assignment experiments.

CTT peptides were purified using Glutathione Sepharose Fast Flow in PBS following the protocol directions provided by the manufacturer (Cytiva). After purification, the peptides were desalted into BRB20 (20 mM PIPES, 1 mM EGTA, 1 mM $MgCl_2$ pH 7) except as noted and the GST tag was cleaved using thrombin that had been desalted into BRB20 pH 7. The cleavage proceeded for 16 hours at room temperature on a nutator. After cleavage, the sample was boiled for 10 minutes to precipitate out the cleaved GST, thrombin, and any remaining uncleaved CTT peptide. The boiled sample was spun at 16,000 x g for 30 minutes and the peptide was collected in the supernatant. It was filtered using a 0.1 um membrane. The protein concentration was determined using BCA (Thermo Scientific).

CAP-Gly domain expression and purification. Human CLIP170 subdomain CAP-Gly2 (156-279) was heterologously expressed in *E. coli* BL21 RIL cells using pET 28 which includes a His purification tag. Unlabeled cultures were grown to OD 0.6-0.8 in LB and induced at 23C for 16 hours with 1 mM IPTG. Labeled cultures for NMR were grown in LB to OD 0.6-0.8, washed in M9 salts (6.2 g/L $Na_2HPO_4$, 3 g/L $KH_2PO_4$, 0.5 g/L NaCl pH 7.4), resuspended in M9 media containing $^{15}NH_4Cl$, and induced at 23C for 16 hours with 1 mM IPTG.

The CAP-Gly2 domain was purified using TALON resin (Takara Bio). All buffers used for purification had a base composition of 25 mM Tris, 500 mM NaCl, 1 mM PMSF pH 8.0. The column was equilibrated in 5 CV of base buffer with 5 mM Imidazole. After a 20-minute incubation with the resin, the clarified lysate was washed through the column with 10 CV of the equilibration buffer followed by 15 CV of base buffer containing 10 mM Imidazole. Finally, the immobilized protein was eluted using 3 CV of base buffer containing 200 mM Imidazole. The eluted protein was concentrated using a 3k concentrator and dialyzed with three, one-liter changes into BRB20 pH 7.

NMR experiments

NMR experiments were performed using an Agilent 600MHz INOVA spectrometer using standard BioPack experiments. For peptide assignment experiments, nonuniform sampling schedules and SMILE reconstruction were used. Experiments were performed at 10C using a RT probe. Data was analyzed using NMRPipe and CCPN Analysis. Samples were a minimum starting volume of 500 μL for titration experiments 300μL for other experiment and contained 10% D2O for locking the spectrometer and 150 mM TSP for peak position referencing. Thin-walled shigemi tubes were used in all experiments.

Assignment and side-chain experiments were performed on 13C, 15N samples, with other experiments performed on 15N samples unless noted. Assignment experiments were Varian Biopack experiments, typically: HNCO, HN(CA)CO, HNCACB, either CBCA(CO)NH or CC(CO)NH, and occasionally HNN.

For the pH titration experiments, samples were started at pH 7 and then brought down to pH 3 in 0.25 pH unit increments using HCl and KOH. The added volumes of acid or base was tracked and in total did not exceed 20% of the sample volume. A micro pH probe was used.

To monitor Asp and Glu pKa values directly, the 13C-detect CBCGCO pulse sequence was used (9). This experiment correlates the aliphatic Asp/Glu side chain carbon with the side chain carboxylic acid carbon.

For the NMR binding experiments TUBA1A was isotopically labeled with 15NH4Cl and brought to a concentration of either 100 mM or 50 mM and combined with 300 mM of CAP-Gly1 (57-130) or 150 mM CAP-Gly2 (156-279), respectively. All binding experiments were performed in BRB20 at either pH 7 or 6 as indicated and HSQCs were collected.

NMR chemical shift values were obtained by peak-picking in CCPNMR analysis or assign. Data were then exported to MATLAB for fitting.

CD spectroscopy

In the last step of the purification protocol detailed above, the peptides were desalted into 100 mM ammonium sulfate, 10 mM monopotassium phosphate. CD was performed using the Applied Photophysics ChirascanPlus (Leatherhead, UK). Peptides at pH 7, 6, 5.5, 5, 4.5, 4, 3.5, and 3 were measured at wavelengths 188-260 (TUBB, TUBA1A) or 189.5-206 (TUBB3) and data was analyzed after buffer subtraction. Data were deconvolved using SESCA (DS-B4R1 basis set) (10).

Fluorescence anisotropy

FITC labeled TUBA1A peptide was obtained from Genscript. Measurements were carried out using a Tecan Spark plate reader. For the FA assay to determine the $k_D$ of the CAP-Gly2 and TUBA1A interaction, fluorescently labeled TUBA1A was held constant at 10 nM and CAP-Gly2 ranged from 100 mM to 12 nM in two-fold dilutions. All experiments were performed in BRB20 at pH 7 and 6. All experiments were performed in BRB20 at pH 7 and 6. Curves were plotted and fitted in Prism.

For the unlabeled TUBA1A competition assays, CAP-Gly2 was held constant at 6 uM for the experiment at pH 7 and 0.6 uM for the experiment at pH 6 (these were twice the concentrations of the measured $k_D$s from the labeled FA experiment). The labeled TUBA1A was held constant at 10 nM and the unlabeled TUBA1A was varied from 1.5 mM to 12 nM in two-fold dilutions. All experiments were performed in BRB20 at pH 7 and 6.

MD simulations

Tubulin C-terminal tail sequences were acquired using HPEPDOCK (11) and AlphaFold3 (12). Each C-terminal tail simulation was then prepared using the standard workflow in the phbuilder tool (13) within the GROMACS constant pH package (13). The N-terminus and C-terminus were made to be neutral (NH2) and negatively charged (COO-), respectively, to more accurately represent being attached to the tubulin body. Each system was then solvated using TIP3 water with $Na^+$ and $Cl^-$ added to both neutralize the system charge and set the ionic strength to 50 mM. Simulations were carried out using GROMACS constant pH using the CHARMM36 force field (14). Following heating and equilibrating for 10 ps each, each system was simulated for 1 μs at 300°K in an NpT ensemble with 1 atm pressure. We utilized the *genparams* tool from phbuilder to assign values for the molecular dynamics parameters (.mdp) file, adjusting the energy calculations and trajectory write out to every 1ps using 2fs timesteps. We performed 3 independent 1 μs simulations at pH 6.0 for

TUBA1A and TUBB. In addition, we performed 1 μs simulations for TUBA1A at pH 4.5, 5.0, 5.5, 6.5, and 7.0, along with 2 independent 1 μs simulations of Tub2 (yeast) at pH 6.0. Poisson-Boltzmann calculations of residue pKa were determined using pypka (15). Other analysis was done using bio3D (16) and R, and images were created using VMD(17).

Cin8 microtubule binding assay

One-liter cultures of yeast cells expressing Cin8-3GFP and wild-type Tub2/β-tubulin, Cin8-3GFP and tub2-430Δ that lacks β-CTT, or an extra copy of a fluorescently tagged GFP-Tub1/α-tubulin were grown to log phase in rich media, pelleted at 4,000 g at 4°C, and then washed twice with cold $H_2O$. The cell pellet was resuspended in a minimal volume of 10X BRB80 (80 mM PIPES, 1 mM $MgCl_2$, 1 mM EGTA) plus 150 mM KCl, snap frozen in liquid nitrogen and stored at -80°C. Frozen cells were lysed in 2 x 110 second cycles of 30 Hz milling in a chilled 50 ml grinding jar on a Mixer Mill MM 400 (Retsch, Hamburg, Germany). For each imaging experiment, two aliquots of approximately 100mg of yeast lysate powder were added to prechilled tubes. The "polymerization sample" was reconstituted in 0.1 μl of 10X BRB80 plus 150 mM KCl per mg of powdered lysate, and 1 μl of yeast protease inhibitor cocktail (Sigma-Aldrich P8215) was added. The "imaging sample" was reconstituted using a ratio of 2 μl of 10X BRB80 plus 150 mM KCl per mg of powdered lysate, and 1 μl of yeast protease inhibitor cocktail (Sigma-Aldrich P8215) was added. Both samples were briefly vortexed, thawed on ice for 10 minutes, and clarified by centrifugation at 100,000 g for 30 minutes at 4°C. Supernatants were transferred to chilled microcentrifuge tube. The polymerization sample was approximately 100 mg/ml protein, and the imaging sample was approximately 50 mg/ml.

Reaction chambers were assembled with a plasma cleaned and HMDS silanized (12) 22 $mm^2$ coverslip and an 18-by-18 mm coverslip separated by single ply strips of parafilm in a custom-made stage insert. Anti-rhodamine antibody (Thermo Fisher A-6397) was diluted to 10 μg/mL in cold BRB80, added to the chamber, incubated for 5 min at room temperature, and then the chamber was then washed with 100 μl of BRB80. 50 μl of 1% pluornic-F127 in BRB80 was added to the chamber, incubated for 5 min at room temperature, and then washed with 100 μl of BRB80. GMPCPP-stabilized porcine brain microtubule seeds containing 23% rhodamine tubulin (Cytosekeleton, Inc TL590M) were then added to the chamber, incubated for 30 sec, and then washed with 200 μl of BRB80.

To polymerize microtubules, a 50 μl reaction consisting of 10 μl of the polymerization sample (approximately 20 mg/ml final concentration) in BRB80 pH6.9 containing 1 mM $MgCl_2$, 8 mM ADP, 1 mM GTP, and 2 μM epothilone A was added to a prepared chamber and incubated for 20 min at 30°C. Polymerization was confirmed by TIRF microscopy, based on imaging Cin8-3GFP signal decorating unlabeled microtubules. The chamber was then washed twice with 50 μl of 750 mM KCl, 25 μM epothilone A in BRB80 for 3 minutes each, and then once for 3 minutes with 100 μl of BRB80. A 500 μl imaging reaction was then prepared with 50 μl of the imaging samples (approximately 5 mg/ml final concentration) in BRB80 pH5.7 containing 1 mM $MgCl_2$, 2 mM ATP, 1 mM GTP, and 25 μM of epothilone A. 50 μl of this reaction was flowed into the chamber and incubated at 30°C for 4 minutes prior to imaging. After imaging at a particular pH, the imaging sample pH was increased by ~0.2 units using KOH, 50 μl was flowed into the chamber, incubated 4 minutes at 30ºC, and imaged.

TIRF images were collected on a Nikon Ti-E microscope equipped with a 1.49 NA 100× CFI160 Apochromat objective, TIRF illuminator, OBIS 488-nm and Sapphire 561-nm lasers (Coherent, Santa Clara, CA), W-View GEMINI image splitting optics (Hamamatsu Photonics, Hamamatsu, Japan) and an ORCA-Flash 4.0 LT sCMOS camera (Hammamatsu Photonics). Stage was heated to 30°C using an ASI 400 Air Stream Incubator (NEVTEK,

Williamsville, VA). At least 10 images of different fields were collected at each pH step, for each sample.

To quantify fluorescent intensity, values were recorded from a 3-pixel wide line drawn along the microtubule. For each field of view a 13-by-13 pixel box was drawn and used for background subtraction. Sum intensity was determined as the background-subtracted sum of all values along the microtubule. To quantify Cin8-3GFP foci, the "findpeaks" function in MATLAB (MathWorks, Natick, MA) was used to objectively determine foci, and the number and background-subtracted intensity of each foci was recorded.

### SI References


1. P. Findeisen, *et al.*, Six subgroups and extensive recent duplications characterize the evolution of the eukaryotic tubulin protein family. *Genome Biol Evol* **6**, 2274–2288 (2014).

2. D. A. Opulente, *et al.*, Genomic factors shape carbon and nitrogen metabolic niche breadth across Saccharomycotina yeasts. *Science* **384**, eadj4503 (2024).

3. V. Redeker, J. Rossier, A. Frankfurter, Posttranslational modifications of the C-terminus of alpha-tubulin in adult rat brain: alpha 4 is glutamylated at two residues. *Biochemistry* **37**, 14838–14844 (1998).

4. J. Mary, V. Redeker, J. P. Le Caer, J. C. Promé, J. Rossier, Class I and IVa beta-tubulin isotypes expressed in adult mouse brain are glutamylated. *FEBS Lett* **353**, 89–94 (1994).

5. A. Rudiger, M. Rudiger, K. Weber, D. Schomburg, Characterization of Posttranslational Modifications of Brain Tubulin by Matrix-Assisted Laser Desorption/Ionization Mass Spectrometry: Direct One-Step Analysis of a Limited Subtilisin Digest. *Analytical Biochemistry* **224**, 532–537 (1995).

6. J. E. Alexander, *et al.*, Characterization of posttranslational modifications in neuron-specific class III beta-tubulin by mass spectrometry. *Proc Natl Acad Sci U S A* **88**, 4685–4689 (1991).

7. B. Eddé, *et al.*, Posttranslational glutamylation of alpha-tubulin. *Science* **247**, 83–85 (1990).

8. V. Redeker, R. Melki, D. Promé, J. P. Le Caer, J. Rossier, Structure of tubulin C-terminal domain obtained by subtilisin treatment. The major alpha and beta tubulin isotypes from pig brain are glutamylated. *FEBS Lett* **313**, 185–192 (1992).

9. C. A. Castañeda, *et al.*, Molecular determinants of the pKa values of Asp and Glu residues in staphylococcal nuclease. *Proteins: Structure, Function, and Bioinformatics* **77**, 570–588 (2009).

10. G. Nagy, M. Igaev, N. C. Jones, S. V. Hoffmann, H. Grubmüller, SESCA: Predicting Circular Dichroism Spectra from Protein Molecular Structures. *J. Chem. Theory Comput.* **15**, 5087–5102 (2019).

11. P. Zhou, B. Jin, H. Li, S.-Y. Huang, HPEPDOCK: a web server for blind peptide–protein docking based on a hierarchical algorithm. *Nucleic Acids Research* **46**, W443–W450 (2018).

12. J. Abramson, *et al.*, Accurate structure prediction of biomolecular interactions with AlphaFold 3. *Nature* **630**, 493–500 (2024).

13. A. Jansen, N. Aho, G. Groenhof, P. Buslaev, B. Hess, phbuilder: A Tool for Efficiently Setting up Constant pH Molecular Dynamics Simulations in GROMACS. *J. Chem. Inf. Model.* **64**, 567–574 (2024).

14. R. B. Best, *et al.*, Optimization of the Additive CHARMM All-Atom Protein Force Field Targeting Improved Sampling of the Backbone ϕ, ψ and Side-Chain χ1 and χ2 Dihedral Angles. *J. Chem. Theory Comput.* **8**, 3257–3273 (2012).

15. P. B. P. S. Reis, D. Vila-Viçosa, W. Rocchia, M. Machuqueiro, PypKa: A Flexible Python Module for Poisson–Boltzmann-Based pKa Calculations. *J. Chem. Inf. Model.* **60**, 4442–4448 (2020).

16. B. J. Grant, A. P. C. Rodrigues, K. M. ElSawy, J. A. McCammon, L. S. D. Caves, Bio3d: an R package for the comparative analysis of protein structures. *Bioinformatics* **22**, 2695–2696 (2006).

17. W. Humphrey, A. Dalke, K. Schulten, VMD: Visual molecular dynamics. *Journal of Molecular Graphics* **14**, 33–38 (1996).
